# Asymmetric subspecies-selection of an NLR-TF immune module and reconstruction of broad-spectrum disease resistance in rice

**DOI:** 10.1101/2025.02.06.636770

**Authors:** Hui Lin, Fudan Chen, Guanyun Cheng, Bingxiao Yan, Meng Yuan, Jie Qiu, Ying Chen, Yijie Wang, Kaixuan Cui, Xiangyu Gong, Shasha Liu, Jiyun Liu, Jianjun Wang, Rongbai Li, Bizeng Mao, Jianlong Xu, Jong-Seong Jeon, Xuehui Huang, Bin Han, Yiwen Deng, Gongyou Chen, Zuhua He

## Abstract

Artificial selection has greatly shaped crop agronomic traits; however, the mechanistic basis of immunity selection has remained elusive. This study identifies a new rice NLR XA48 and its downstream transcription factor OsVOZ1, which confer bacterial blight resistance. XA48 perceives an ancient pathogen effector, XopG, to activates effector-triggered immunity (ETI). The XA48-OsVOZ1 module has undergone subspecies-specific selection. *Xa48* is retained in *indica* but functionally lost in *japonica* rice. OsVOZ1 has also diverged into two haplotypes, *indica* kept both OsVOZ1^A/S^ alleles that match XA48; while *japonica* only inherited OsVOZ1^A^ that greatly decreases yield when *Xa48* is reintroduced into *japonica*, mechanistically explaining the *Xa48* loss in *japonica*. We resurrected wild rice broad-spectrum resistance by stacking XA48-mediated ETI with XA21-mediated pattern-triggered immunity (PTI). Thus, our study reveals that the asymmetric selection of an NLR-TF module shapes both disease resistance and reproduction, and provides a paradigm for breeding crops by harnessing the immunity of wild relatives.

---

Plants have developed a sophisticated immune system to combat pathogens: PTI and ETI, which are governed by cell surface pattern-recognition receptors (PRRs) and intracellular NLR receptors, respecively^1–3^. Plant immunity has been shaped by crop domestication and breeding practices that aim to balance tradeoffs between growth and defense under diverse agricultural conditions^4–6^. It is unclear whether plant *R* genes and their associated immune components have been artificially selected during crop domestication^7,8^. *Xanthomonas oryzae* pv. *oryzae* (*Xoo*) causes bacterial leaf blight (BLB), one of the most devastating diseases of rice, and BLB outbreaks frequently occur after typhoons and heavy rainstorms^9,10^. The first cloned *Xa* gene, *Xa21*, originated from the wild rice species *Oryza longistaminata*, which encodes a RLK that triggers race-specific PTI via perceiving the PAMP RaxX^11,12^. However, *Xa21* does not impart disease resistance to *Xoo* strains from Northeast Asia due to the evolution of virulence in *Xoo*^13,14^. It is particularly important to isolate *an Xa* gene that can complement *Xa21* for breeding modern rice with broad-spectrum resistance (BSR) to *Xoo* in face of the ever changing climate. Intriguingly, most functional *Xa* genes/loci have been isolated from wild relatives or represent recessive *xa* alleles of susceptibility genes^15^, suggesting negative selection of *Xoo R* genes during rice domestication. Given that extensive artificial selection has typically targeted high yielding and reliable disease resistance during breeding^6,16^, answering the question of whether and how *R* gene selection has occurred is particularly important for crop breeding, and we also need a practical breeding example to build BSR through integrating PTI and ETI in a real crop.

## Results

### Functional identification of the NLR gene *Xa48* that confers lifetime *Xoo* resistance

We combined map-based cloning and genome-wide association study (GWAS) to isolate an *Xa* gene that confers *Xoo* resistance and complements *Xa21*. We first identified an *indica* variety Shuangkezao (SKZ) that exhibits strong resistance to the Korean *Xoo* strain J18 (DY89031), but high susceptibility to the Philippine *Xoo* strain PXO99A; while *japonica* TP309 line 106 harboring *Xa21* is reverse^13^ (Extended Data Fig. 1a), suggesting that SKZ has a *R* gene that confers race-specific resistance to *Xoo* and complements *Xa21*. The gene, designated *Xa48*, was identified by map-based cloning using a cross between SKZ and the *japonica* variety Nipponbare (NIPB). *Xa48* was mapped to a 65-kb region on chromosome 3 using a bacterial artificial chromosome (BAC) of SKZ DNA (Extended Data Fig. 1b). Sequencing of the BAC clone revealed four candidates (*C1* to *C4*) harboring deletion/insertion polymorphisms in the NIPB and SKZ genomes (Fig. 1a).

**Fig. 1.**
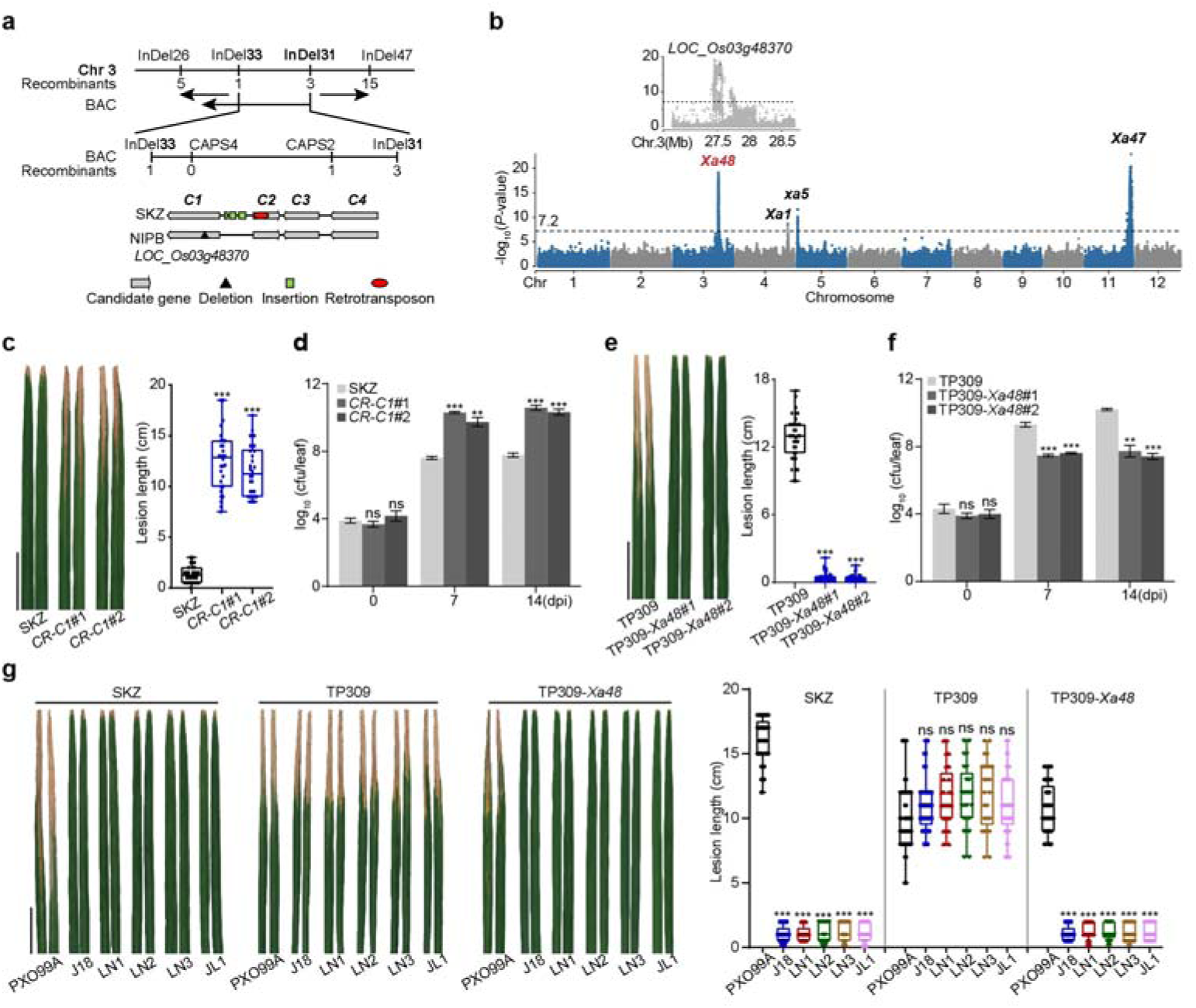
***Xa48* confers lifetime resistance against northeast Asian bacterial blight strains. a**, *Xa48* was preliminarily mapped to chromosome 3 between two insertion/deletion (InDel) markers InDel31 and InDel33, and was finely mapped to a 65-kb interval on a BAC of SKZ. Sequence analysis of the BAC revealed four candidates (*C1-C4*) located in the mapping region. A deletion and SNPs were found within candidate gene *C1* (*LOC_Os03g48370*) in NIPB. **b**, Manhattan plot of GWAS for resistance to *Xoo* strain J18 in rice accessions. The genome-wide significant value threshold (6.35E-8) is indicated by a horizontal dash-dot line. A zoomed-in plot for the GWAS peak was shown with two significantly associated nonsynonymous SNPs in the *Xa48* region highlighted. **c**, **d**, *Xa48* knockout *CR-C1* plants completely lost resistance to strain J18, the lesion length was measured at 14 days post-inoculation (dpi) (**c**). Bacterial populations per leaf were measured at 0, 7 and 14 dpi (**d**). **e**, **f**, Complement transgenic plants (TP309-*Xa48*) exhibited high *Xoo* resistance. The lesion length at 14 dpi (**e**) and bacterial populations at 0, 7 and 14 dpi (**f**) were measured. **g**, *Xa48* also confers resistance to other northeast Asian strains. Plants were inoculated with *Xoo* strain JL1, LN1, LN2 and LN3 collected from Northeast China, with J18 and PXO99A as avirulent and virulent control strain, respectively. Disease symptom was measured at 14 dpi. Data were shown as mean ± SD, n ≥ 25 (**c**, **e**, **g**) and n = 3 (**d**, **f**). Scale bars, 5 cm (**c**, **e**, **g**). For **c** to **g**, asterisks represented statistical significance (\*\**P* < 0.01, \*\*\**P* < 0.001, two-tailed Student’s t-test). ns, not significant. Experiments were independently repeated three times with similar results (**c-g**).

A GWAS analysis for resistance to strain J18 was executed across 1,945 rice accessions and revealed four associated loci on chromosomes 3, 4, 5 and 11 (Fig. 1b). Three of the four loci contained the *Xa* alleles *Xa1*, *xa5* and *Xa47*, previously reported^17–19^. The other locus exhibits a substantial GWAS signature peak that overlaps with the mapped *Xa48* locus. Next, we generated CRISPR/Cas9 knockout mutants (*CR-C1*, *CR-C2*, *CR-C3*, and *CR-C4*) in the four loci, and discovered that *CR-C1* exhibited a susceptible phenotype to J18 (Fig. 1c, d, Extended Data Fig. 1c, d). Compared with SKZ, the *C1* locus in NIPB has a single-base deletion (Extended Data Fig. 1e). These results strongly suggest that *C1* is the candidate gene for *Xa48* (*LOC_Os03g48370*), which encodes an NLR receptor consisting of 1094 amino acids (aa) (Extended Data Fig. 1e). To further confirm *Xa48*, a construct containing *Xa48* genomic DNA and a 3,000-bp promoter region was transferred into *japonica* TP309 and NIPB (Extended Data Fig. 1f). The resulting TP309-*Xa48* and NIPB-*Xa48* lines were highly resistant to J18 (Fig. 1e, f, Extended Data Fig. 1g, h). Similarly, *Xa48* confers resistance to *Xoo* strains from Northeast Asia, including LN2^13^ (Fig. 1g). Therefore, we conclude that *Xa48* is an NLR gene responsible for race-specific resistance to *Xoo* strains originating from Northeast Asia.

In contrast to other *Xa* genes that generally confer resistance only in adult plants^15^, *Xa48* also confers *Xoo* resistance in seedlings (Extended Data Fig. 2a-d), which supports its potential use in rice breeding. *Xa48* is expressed at high levels in leaves and panicles, and it is not induced by pathogen infection (Extended Data Fig. 2e-g). Furthermore, RNA-seq analysis of wild type TP309 and TP309-*Xa48* infected with *Xoo* revealed differentially expressed genes (DEGs) enriched in the biosynthesis of salicylic acid, ethylene and phytoalexin, implying the involvement of these signals in *Xa48*-mediated immunity (Extended Data Fig. 2h, i).

### XA48 perceives cognate effector AvrXa48 (XopG) and activates immunity

Tn*5*-mediated mutagenesis of *Xoo* J18 was unsuccessful, so strain LN2 was used instead to search for *avrXa48*. A LN2 mutant library was generated with Tn*5* and 20,000 insertion mutants were inoculated to *Xa48* rice. Two *Xoo* mutants were obtained that exhibited virulence on *Xa48* rice (Fig. 2a, Extended Data Fig. 3a-c). Plasmid rescue and complementation analysis of the *Xoo* mutants successfully identified *avrXa48*, which is a predicted type III effector annotated as XopG without functional identification. For consistency, we use XopG hereafter. Production of XopG in *Xoo* PXO99A (PXO99A/XopG_LN2_) converted PXO99A into an avirulent strain on *Xa48* rice (Fig. 2b, Extended Data Fig. 3d). Thus, XopG is the cognate effector to XA48.

**Fig. 2.**
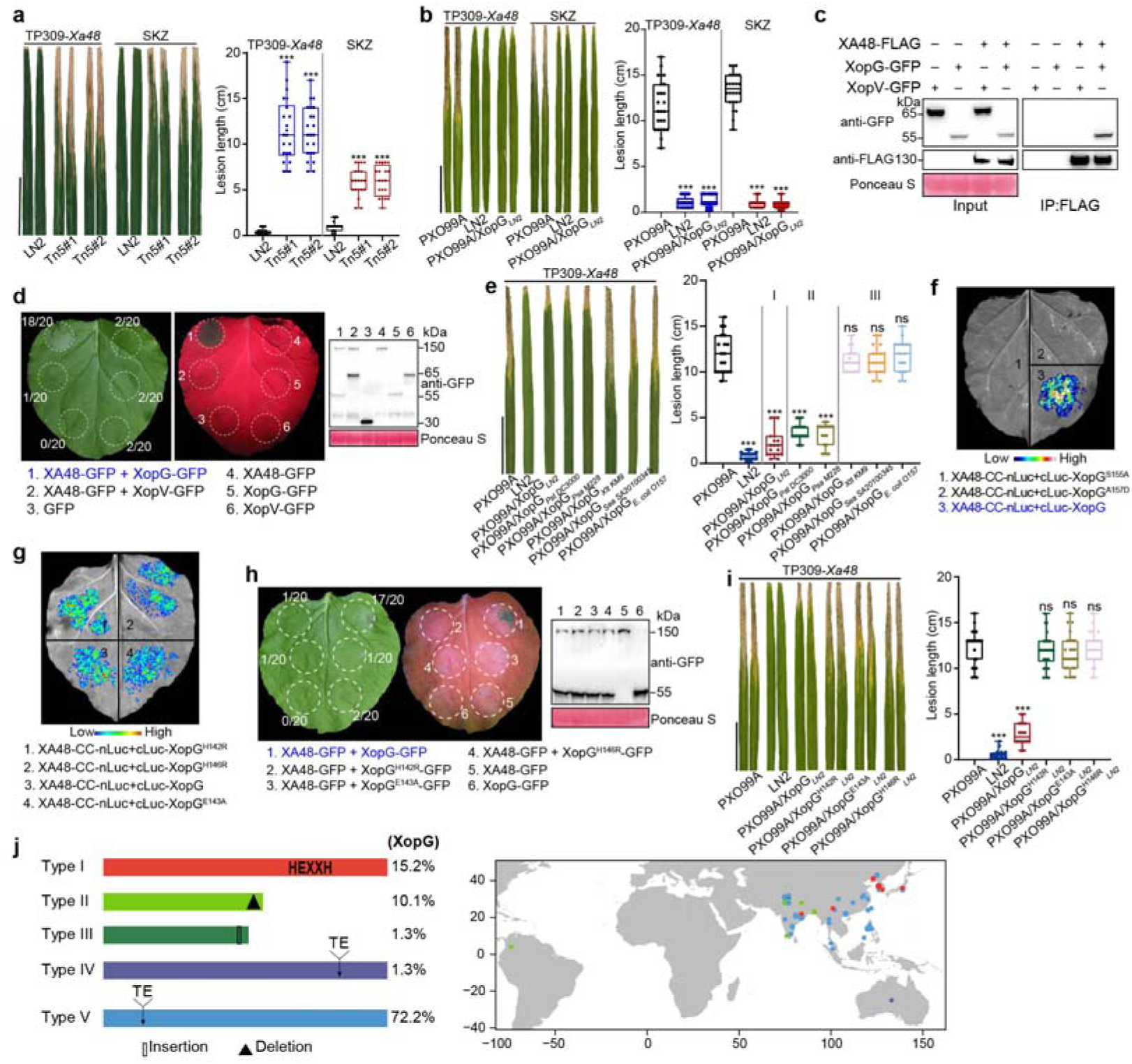
XA48 perceives the cognate effector XopG to activate immunity. **a**, Tn*5* transposon-inserted mutants of strain LN2 lost avirulence to TP309-*Xa48* and SKZ. The lesion length was measured at 14 dpi. **b**, The introduction of XopG*_LN2_* changed PXO99A virulence into avirulence to TP309-*Xa48* and SKZ. **c**, Interaction of XA48-FLAG with XopG-GFP detected by co-IP in transgenic rice plants. Fusion proteins were immunodetected using anti-GFP or anti-FLAG antibody. Another *Xoo* effector protein XopV, which does not interact with XA48, was used as a negative control. **d**, Co-expression of XA48 and XopG induced hypersensitive response cell death in *N. benthamiana* leaves. Protein expression was detected by Western blot. Numbers in parentheses indicate the leaves that exhibited cell-death symptoms. **e**, XopG-like proteins from *Xoo* strain LN2, *Pseudomonas syringae* pv. *tomato* DC3000 (*Pst*), P. *syringae* pv. *actinidae* M228 (*Psa*), *Xanthomonas translucens* pv. *translucens* Km9 (*Xtt*), *Salmonella enterica* subsp. *arizonae* SA20100345 (*Sea*) and *Escherichia coli* O157, were individually introduced into strain PXO99A. Note that Clade I and II XopG-like proteins were avirulent to *Xa48*. LN2 and PXO99A-XopG_LN2_ were used as avirulent controls. **f**, Mutants of XopG^S155A^ and XopG^A157D^ lost interaction with XA48-CC, as detected by SLC. **g**, **h**, XopG mutants (H142R, E143R, H146R) interact with XA48-CC, detected by split luciferase complementation (SLC) (**g**), but could not trigger XA48-mediaed cell death in *N. benthamiana* leaves (**h**). Protein expression was detected by Western blot. Numbers in parentheses indicate leaves exhibited cell-death symptoms. **i**, Disease resistance phenotype and lesion length of TP309-*Xa48* inoculated with PXO99A/XopG*_LN2_* and PXO99A expressing the three HEXXH motif mutant variants (PXO99A/XopG*_LN2_* ^H142R^, PXO99A/XopG*_LN2_* ^E143R^, and PXO99A/XopG*_LN2_* ^H146R^). Note that the HEXXH motif is critical to its function in *Xa48*-mediated resistance. LN2 and PXO99A-XopG*_LN2_* were used as avirulent controls. **j**, XopG homologs among a total of 76 *Xanthomonas* are classified into five types based on protein sequences. Only Type I contains full-length XopG, whereas other types are truncated. TE, transposable element. The geographical distribution of the XopG homologs reveals that functional XopG in Type I, is mainly found in strains from northeast Asia. Data were shown as mean ± SD, n ≥ 20, asterisks represented statistical significance (\*\*\**P* < 0.001, two-tailed Student’s t-test). ns, not significant. Scale bars, 5 cm (**a, b, e, i**). Experiments were independently repeated twice with similar results (**a-i**).

To explore the function of XopG in XA48-mediated ETI, we investigated whether XA48 directly interacts with XopG. An interaction between XopG and the coiled-coil (CC) domain of XA48 was observed in split luciferase complementation (SLC), bimolecular fluorescence complementation (BiFC) and co-immunoprecipitation (co-IP) assays (Fig. 2c, Extended Data Fig. 3e, f). Furthermore, we established that the co-expression of XA48 and XopG induced cell death in *Nicotiana benthamiana* (Fig. 2d), and XopG induced the oligomerization of XA48 (Extended Data Fig. 3g). The effector-induced NLR oligomerization was also observed for the TNL receptor ROQ1 by another *Xanthomonas* effector XopQ in *Arabidopsis*^20^. Therefore, our findings show XA48 perceives XopG to activate immunity.

### XopG is a conserved effector in bacterial pathogens

XopG contains a peptidase domain M91 with a conserved HEXXH motif (Extended Data Fig. 3h). This effector family includes the *E. coli* effector protein NleD, which cleaves and inactivates c-Jun N-terminal kinase^21^. To test the endopeptidase activity of XopG, self-cleavage of purified recombinant XopG fused to the maltose-binding protein (XopG-MBP) was observed *in vitro* (Extended Data Fig. 3i); however, there was no evidence that XopG can cleave XA48 *in vivo* (Extended Data Fig. 3j), which supported our observation that XopG promotes XA48 oligomerization (Extended Data Fig. 3g). Additionally, XopG was localized in the plasma membrane and nucleus, and it exhibited co-localization with XA48 (Extended Data Fig. 3k).

We analyzed 38 proteins orthologous to XopG from other pathogenic bacteria and found they could be divided into three clades (Extended Data Fig. 4a). The *xopG* homolog genes were transformed into *Xoo* PXO99A, and the resulting PXO99A strains were inoculated to *Xa48* rice. The PXO99A strains expressing XopG-like proteins from Clades I and II were avirulent on *Xa48* plants, whereas those expressing Clade III proteins remained virulent (Fig. 2e, Extended Data Fig. 4b); this indicated that the XopG-like proteins from Clades I and II are likely to be recognized by XA48 to activate immunity. Next, an in-depth analysis revealed that the S and A aa residues at position 155 and 157 are conserved in XopG like proteins in Clades I and II but not in Clade III (Extended Data Fig. 4a). Further, site-directed mutagenesis was used to mutate S155 to A and A157 to D; the resulting XopG^S155A^ and XopG^A157D^ variants failed to interact with XA48-CC (Fig. 2f). When the two XopG variants were expressed in strain PXO99A (PXO99A/XopG^S155A^ and PXO99A/XopG^A157D^) and inoculated to *Xa48* plants, the strains lost the ability to trigger XA48-mediated resistance (Extended Data Fig. 4c). We suspected that XopG belongs to an ancient effector family that impairs basal resistance to *Xoo* in rice (Extended Data Fig. 5a), and we speculated that the conserved HEXXH motif plays a critical role in XopG function. This hypothesis was tested by mutating 142H to R, 143E to A, and 146H to R. The H142R, E143A, and H146R mutants interacted with XA48-CC but could not induce XA48-mediated cell death (Fig. 2g, h). Furthermore, these mutants also lost the ability to trigger XA48*-*mediated resistance in rice (Fig. 2i, Extended Data Fig. 5b).

The sequence variability among XopG orthologs was investigated in 76 *Xoo* stains using available genome sequences (Supplementary Table 1), and the orthologs were classified into five structural types. Type I contained full-length XopG, whereas other types either lacked the HEXXH motif or were truncated (Fig. 2j). Consistent with the expected function of the HEXXH motif, *Xoo* strains LN2, LN4 and LN18 harbor a Type I XopG and are avirulent to *Xa48* rice, whereas virulent strains PXO99A and JL28 harbor a Type V XopG (Extended Data Fig. 5c, d). The geographical distribution of *Xoo* strains revealed that Type I strains originated from Northeast Asia, whereas *Xoo* strains with mutated forms of XopG were mainly distributed in South Asia. These results support our previous finding that *Xoo* effectors exhibit a high level of divergence in both sequence and pathogenicity^13^. We propose that the prolonged arms race between XA48 and XopG has likely exerted a selective pressure on the evolution of *Xa48* (see below for further details).

### XA48 interacts with OsVOZ transcription factors for immune signaling

To elucidate the underlying molecular mechanism for XA48-mediated resistance, we conducted a yeast two-hybrid (Y2H) screen^22^ to isolate XA48-interacting proteins.

The transcription factors Vascular PLANT ONE-ZINC FINGER proteins, OsVOZ1 and OsVOZ2, were identified (Fig. 3a, b, Extended Data Fig. 6a). The XA48-OsVOZ1/2 interaction was validated *in planta* using SLC and co-IP assays (Fig. 3c-f, Extended Data Fig. 6b), and the results showed that OsVOZ1/2 interacted with full-length XA48 and its CC domain. We transiently co-expressed OsVOZ1/OsVOZ2-mCherry with XA48-YFP, and observed co-localization of XA48 and OsVOZ1/2 in the plasma membrane and nucleus (Extended Data Fig. 6c, d). This co-localization pattern is similar to that observed in the XA48-XopG association, hinting that XA48 might regulate OsVOZ1 and OsVOZ2.

**Fig. 3.**
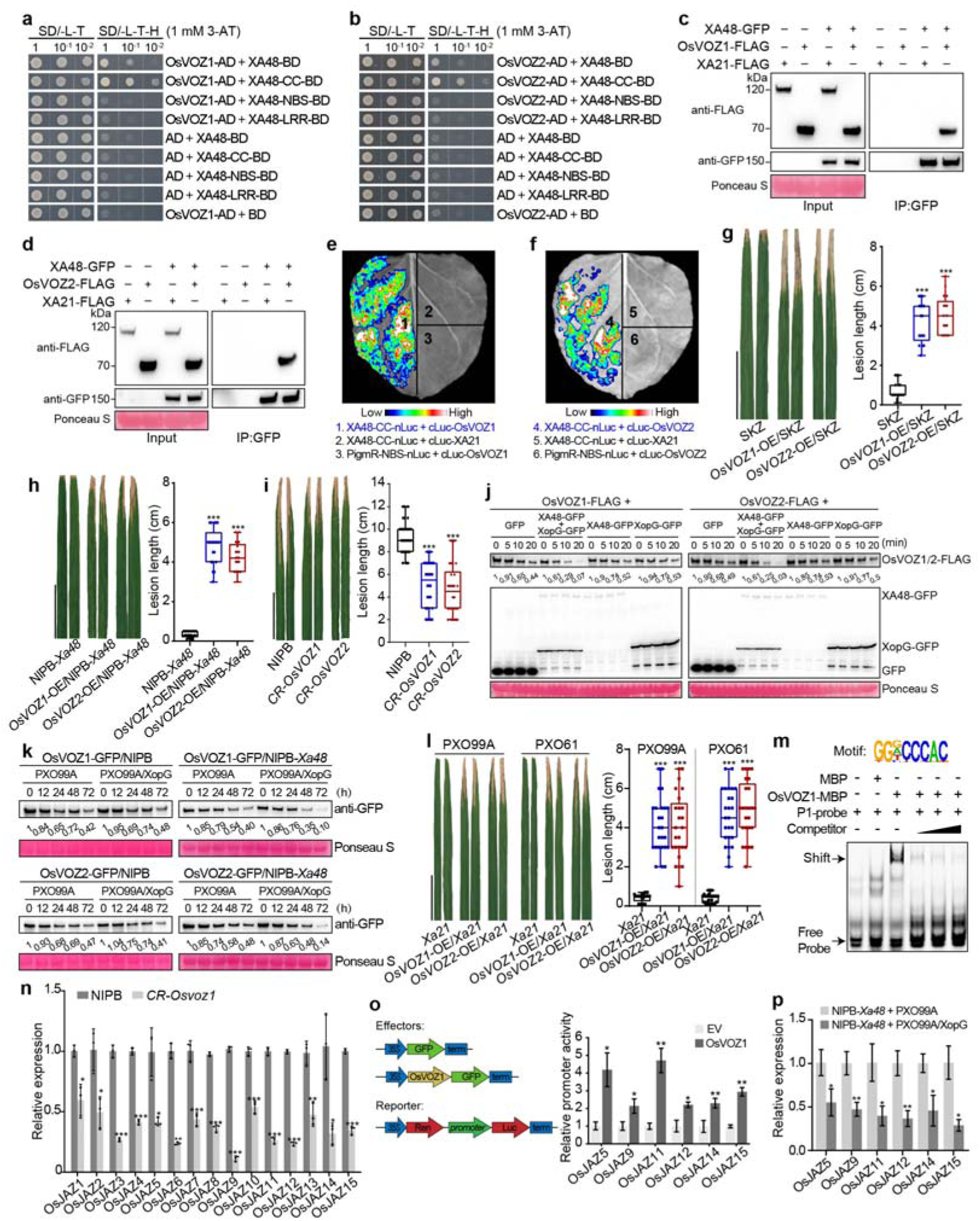
XA48 interacts with and facilitates degradation of the transcription factors OsVOZ1 and OsVOZ2 for immune activation. **a**, **b**, The interaction between XA48 with OsVOZ1 (**a**) and OsVOZ2 (**b**) was detected by yeast two-hybrid (Y2H). Note that XA48 interacts with OsVOZ1/2 through the CC domain. SD/-L-T-H, synthetic dropout (SD) media lacking leucine, tryptophan, and histidine; SD/-L-T, SD media lacking leucine and tryptophan; 3AT, 3-aminotriazole. **c**, **d**, Co-IP assay confirms the interaction of XA48 and OsVOZ1/2. XA21 was expressed as negative control (Pruitt et al., 2015). **e**, **f**, SLC shows the interaction of XA48-CC and OsVOZ1/2 *in planta*. XA21 and PigmR-NBS (Zhai et al., 2022), which do not interact OsVOZs, were used as negative controls. **g**, **h**, OsVOZs negatively regulated XA48-mediated resistance against *Xoo*. Overexpression of *OsVOZ1/2* resulted in decreased J18 resistance in SKZ (**g**) and NIPB-*Xa48* transgenic plants (**h**). **i**, OsVOZs negatively regulate basal defense against *Xoo*. Knockout mutants of *OsVOZ1/2* in NIPB increased resistance to J18. **j**, XA48-XopG promoted degradation of OsVOZ1 (left) and OsVOZ2 (right) in a cell-free assay. Extracts from OsVOZs-FLAG transgenic rice plants were incubated with cell extracts from transgenic rice XA48-GFP and XopG-GFP, and samples were collected over a time course of 0-20 min after mixing. GFP expression alone served as negative controls. **k**, Degradation promotion of OsVOZ1/2 by XopG-XA48 recognition was detected in planta. *OsVOZ*s-GFP/NIPB and *OsVOZ*s-GFP/NIPB-*Xa48* plants were inoculated with PXO99A and PXO99A/XopG, respectively. proteins were collected over a time course of 0-72 hours post inoculation (hpi) for Western blotting. **l**, OsVOZs negatively regulates *Xa21*-mediated resistance. Overexpression of *OsVOZ1/2* resulted into decreased resistance to PXO99A and PXO61 in *OsVOZs*-OE/*Xa21* transgenic plants. **m**, Binding specificity of OsVOZ1 towards the CCCAC motif of the *JAZ* promoter. Biotin-labeled probes were incubated with MBP or OsVOZ1-MBP, with unlabeled competitor fragments co-incubated. The bands representing the DNA-protein complexes (shift) and the free probes are indicated by arrows. **n**, *OsJAZs* showed decreased expression in *CR-Osvoz1* plants compared with wildtype NIPB, as detected by qRT-PCR. **o**, OsVOZ1 enhances promoter activity *OsJAZs*. OsVOZ1-GFP activates the expression of the pro*OsJAZs*-LUC fusion reporter in *N. benthamiana*. The LUC activity was measured by normalizing to REN signal. **p**, *OsJAZs* were down-regulated by XA48-XopG. NIPB-*Xa48* were inoculated with PXO99A and PXO99A/XopG, respectively, and leaf samples were collected at 24hpi for RNA preparation. Data were shown as mean ± SD, *n* ≥ 15 (**g**-**i**, **l**), n = 3 (**n**-**p**), asterisks represented statistical significance (*P < 0.05, **P < 0.01, \*\*\**P* < 0.001, two-tailed Student’s t-test). Scale bars, 5 cm (**g**-**i**, **l**). Experiments were independently repeated three times with similar results.

OsVOZ2 was reported to affect *Xoo* virulence^23^. To determine the roles of OsVOZs in XA48-mediated resistance, we generated transgenic plants overexpressing *OsVOZ1/2* (*OsVOZ1*-OE and *OsVOZ2*-OE) in SKZ (*Xa48*, *indica*), NIPB-*Xa48* (*japonica*) and the knockout mutants (*CR-Osvoz1*/*2*) by CRISPR/Cas9 editing (Extended Data Fig. 6e, f). A rice *osvoz1/osvoz2* double mutant was not obtained due to its lethality^24^, and the *OsVOZ1/2*-OE line showed attenuated XA48-mediated resistance (Fig. 3g, h). In contrast, *CR-Osvoz* mutants displayed increased basal resistance when compared with wild type NIPB (Fig. 3i). Therefore, OsVOZs function as negative regulators in both basal and XA48-triggered immunity against *Xoo*. The biological importance of the XA48-OsVOZs interaction in XA48-mediated resistance was then investigated. When the accumulation of OsVOZ1-FLAG and OsVOZ2-FLAG was measured in cell-free protein degradation assays, the presence of XA48 and XopG promoted the degradation of OsVOZ1 and OsVOZ2 (Fig. 3j). The XopG-XA48 promoting degradation of OsVOZ1/2 was also investigated *in planta*. Inoculation with PXO99A/XopG but not PXO99A promoted degradation of OsVOZ1/2 in *OsVOZs*-OE/NIPB-*Xa48* rice as compared to *OsVOZs*-OE/NIPB (Fig. 3k). These results confirmed that OsVOZs undergo degradation to release the suppression of immunity during ETI activation.

OsVOZ1 and OsVOZ2 were previously shown to interact with Piz-t, an NLR that confers resistance to the rice blast fungus, *M. oryzae*. OsVOZ1/2 negatively regulated basal defense and positively regulated Piz-t-mediated ETI, presumably by modulating Piz-t expression and protein levels^24^. To evaluate this possibility in the rice-*Xoo* interaction, plants overexpressing *OsVOZs* were generated in the *Xa21* line 106. When these plants were inoculated with the virulent *Xoo* strains PXO99A and PXO61, OsVOZs were shown to negatively regulate XA21-mediated PTI against *Xoo* (Fig. 3l). Therefore, OsVOZs function in both PTI and ETI against *Xoo*, which may adopt differential signaling mechanisms to cope with pathogens with diverse infection lifestyles.

To dissect the immune suppression function of OsVOZs, Cleavage Under Targets & Tagmentation (CUT&Tag) assay was executed in OsVOZ1-GFP/NIPB transgenic plants. DNA motif (CCCAC) had a high binding affinity for OsVOZ1-MBP (Fig. 3m). Next, RNA-seq analysis of *CR-Osvoz1* and wild type NIPB revealed a set of DEGs involved in defense response, such as amino acid biosynthetic, metabolic process, and jasmonic acid (JA) signaling (Extended Data Fig. 7a). Expression level of *JASMONATE-ZIM-DOMAINs* (*OsJAZs*), encoding JA signaling and defense repressors in rice^25,26^, were downregulated in *CR-Osvoz1* rice as compared to NIPB (Fig. 3n) and enriched the CCCAC motif in their promoters, indicating that JAZ-mediated immune suppression was alleviated. EMSA assays indicated that OsVOZ1-MBP bound specifically to CCCAC motifs in promoters of *OsJAZ5*, *OsJAZ9*, *OsJAZ11*, *OsJAZ12*, *OsJAZ14*, and *OsJAZ15* tested (Extended Data Fig. 7b), suggesting that OsVOZ1-regulates expression of these genes. Binding of OsVOZ1 to *OsJAZ* promoters induced *OsJAZ* expression based on luciferase-to-Renilla (LUC/REN) ratio (Fig. 3o); this suggested that OsVOZ1 activated *OsJAZ* expression by directly binding to their promoters. Furthermore, qRT-PCR analysis revealed that *OsJAZs* expression decreased when NIPB-*Xa48* rice was inoculated with *Xoo* PXO99A/XopG as compared to the wild type PXO99A (Fig. 3p). Therefore, JA signaling at least plays a role in the XA48-OsVOZ1 module-mediated *Xoo* resistance. Collectively, these data establish an immune signaling pathway where the XA48-XopG immune complex triggers ETI by interacting with and promoting degradation of OsVOZs thereby inhibiting expression of the JAZ immune suppressors.

### *Xa48* was retained in *indica* but functionally lost in *japonica*

The *Xa48* locus present in *japonica* lines NIPB and TP309 was examined, and both genomes contained a nonfunctional allele due to loss-of-function mutations (Extended Data Fig.1e). Based on this finding, we analyzed the divergence of *Xa48* alleles in the genomes of 3K rice germplasm^27^. A total of 1,945 available accessions were inoculated with *Xoo* strain J18 (Fig. 4a), and 405 accessions (20.8%) were identified with resistance to strain J18. Of these 405 accessions, 145 (35.8%) carried a functional *Xa48*; the others harbored loss-of-function mutations in *Xa48*, suggesting that other *R* genes were active against *Xoo* J18. Intriguingly, rice lines with a functional *Xa48* allele are exclusively *indica* subspecies (Fig. 4a, Supplementary Table 2). This suggests that *Xa48* might have undergo differential selection in the two subspecies, leading to its functional retention in *indica* but loss in *japonica*.

**Fig. 4.**
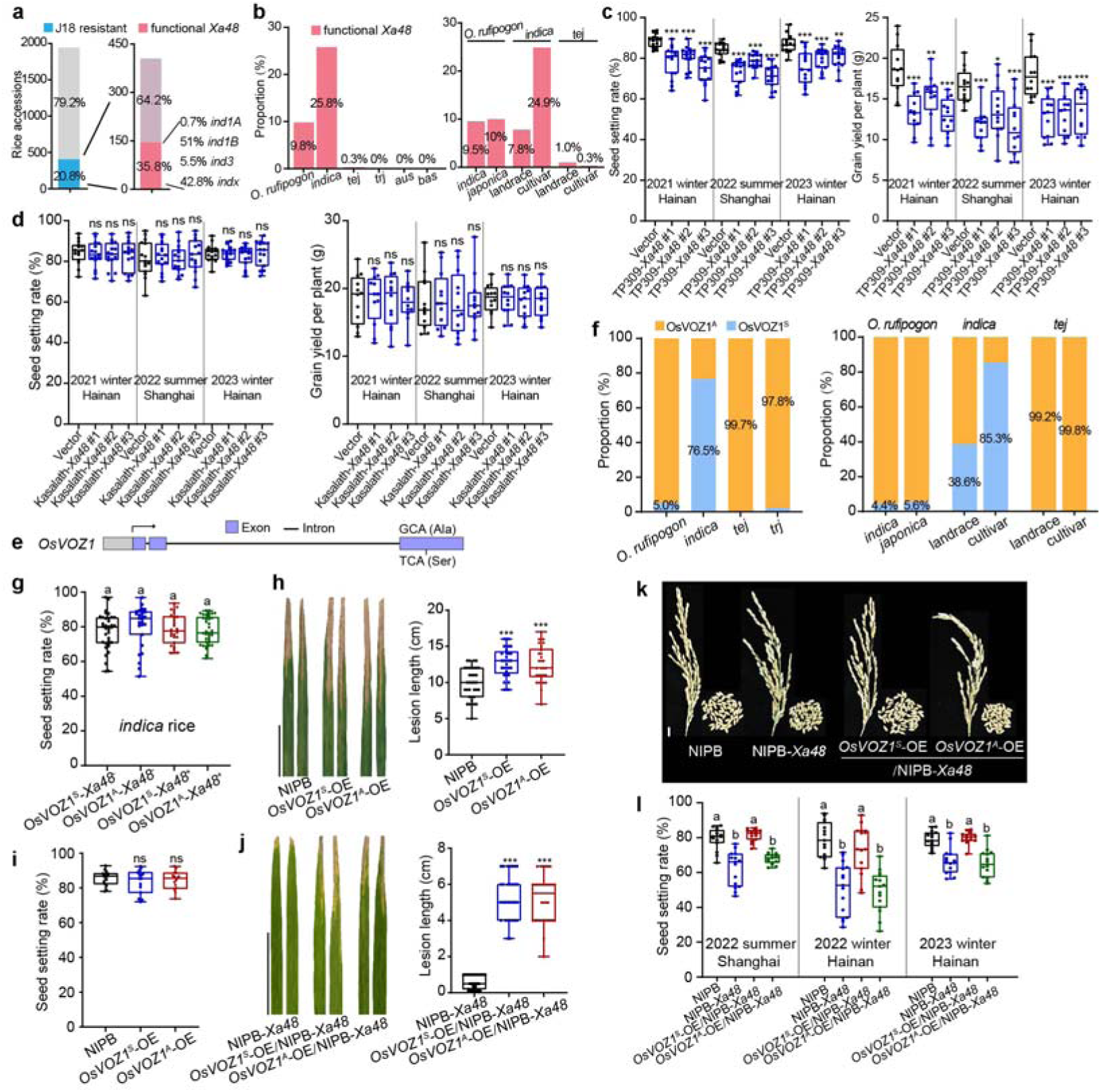
Subspecies-species selection of OsVOZ1 contribute to *Xa48*-induced reproductive penalty in *japonica*. **a**, The frequency of functional *Xa48* was assessed in sequenced rice accessions. *Xa48* was detected in resistant accessions by PCR-based sequencing. Note that near all *Xa48*-containing accessions are *indica* rice, and no *japonica* rice was detected to contain functional but loss-of-function *Xa48* mutants. **b**, Dynamic frequency changes of functional *Xa48* during *indica* and *japonica* domestication and improvement. **c**, The introduction of *Xa48* led to a significantly decrease in grain yield in the *japonica* rice TP309 by reducing seed setting in multiple-locations and seasons field trials. **d**, The introduction of *Xa48* did not affect grain productivity in the *indica* variety Kasalath in multiple-locations and seasons field trials. **e**, *OsVOZ1* exists a major non-synonymous polymorphism SNP^G^ and SNP^T^, resulting in an amino acid change from alanine (OsVOZ1^A^) to serine (OsVOZ1^S^). **f**, Dynamic frequency changes of the OsVOZ1^A^ and OsVOZ1^S^ alleles during *indica* and *japonica* domestication and improvement. Note that nearly all *japonica* accessions carry the OsVOZ1^A^ allele, whereas *indica* rice harbors both alleles. **g**, Seed setting rates of 70 *indica* rice accessions that contain either *OsVOZ1^S^* or *OsVOZ1^A^* haplotype with or without (*Xa48*^+^ or *Xa48*^-^) from the 3K rice genome project, showing no effect of *OsVOZ1^A/S^*alleles on reproductivity in *indica* rice bringing native *Xa48*. **h**, **i**, Overexpression of *OsVOZ1^A^* and *OsVOZ1^S^*alleles significantly decreased resistance to J18 (**h**) without effect on seed setting in NIPB (**i**). Scale bars, 5 cm. **j**, *OsVOZ1^A^* and *OsVOZ1^S^* overexpression compromised *Xa48-*mediated resistance*. OsVOZ1^A^*-OE/NIPB-*Xa48* and *OsVOZ1^S^*-OE/NIPB-*Xa48* were generated by crossing *OsVOZ1^A^*-OE and *OsVOZ1^S^*-OE with NIPB-*Xa48*, respectively. Homozygous plants were selected from the F_2_ populations. Scale bars, 5 cm. **k**, **l**, Overexpression of *OsVOZ1^S^* but not *OsVOZ1^A^* restored normal seed setting in NIPB*-Xa48* (*japonic*a) (**k**). The Field trail of two seasons and locations revealed that the seed setting of *OsVOZ1^S^*-OE/NIPB-*Xa48* was stored as the wild type NIPB (**l**). Scale bars, 1 cm. Data were shown as mean ± SD, *n* ≥ 12 (**c, d, g, i, l**), asterisks represented statistical significance (\**P* < 0.05, \*\**P* < 0.01, \*\*\**P* < 0.001, two-tailed Student’s t-test). ns, not significant. Letters indicate significant differences (*P* < 0.05) determined by two-way analysis of variance (ANOVA) with Tukey’s test (**g**, **l**). Experiments were independently repeated three times with similar results (**c, d, g, I, l**).

Next, we determined the sequence of *Xa48* in other genomic projects with 2,451 Asian rice accessions including wild rice with available genome sequences^28–30^ (Supplementary Table 3). Based on GWAS peak SNPs (*P*=1.47E^-16^), we examined the dynamic frequency change of functional *Xa48* allele during rice domestication and improvement. In the wild rice species, *O. rufipogon*, the functional allele frequency was moderate (9.8%). In *indica*, the allele frequency increased from 7.8% to 24.9% during breeding improvements that occurred from landraces to cultivars. In contrast, the functional allele of temperate *japonica* decreased from a rare frequency in landraces (1.0%) and improved populations (0.3%) (Supplementary Table 4). These results suggest positive and negative selection for *Xa48* in *indica* and *japonica*, respectively. Considering the agroecological-specific distribution of XopG-positive *Xoo* strains in Northeast Asia (Fig. 2j) where *japonica* rice is cultivated, we suggest that a negative selection pressure occurred for XopG in *Xoo* to deal with XA48-mediated resistance.

### *Xa48* decreases grain yield in *japonica*

Our next objective was to determine the underlying mechanism leading to the elimination of *Xa48* in *japonica* but not *indica*. Considering the growth penalty associated with disease resistance in rice^31,32^, agronomic traits were compared in the following rice lines: SKZ vs *CR-Xa48/*SKZ (CRISPR/Cas9 knockout line), Kasalath (without *Xa48*) vs transgenic Kasalath-*Xa48* (*indica*) (Extended Data Fig. 8a), NIPB vs NIPB-*Xa48*, and TP309 vs TP309-*Xa48* (*japonica*). Morphological differences in these transgenic lines were not observed when compared to their respective wild types; however, one exception was the lower plant height observed in NIPB-*Xa48* and TP309-*Xa48* (Extended Data Fig. 8b, c). Interestingly, seed setting rate decreased significantly in *japonica* NIPB-*Xa48* and TP309-*Xa48* but remained unchanged in *indica CR-Xa48/*SKZ and Kasalath-*Xa48* plants based on field trials over multiple seasons and locations, resulting in lower grain yield in NIPB-*Xa48* and TP309-*Xa48* as compared to wild type NIPB and TP309 (Fig. 4c, d, Extended Data Fig. 8d-f). These findings indicated the reintroduction of *Xa48* greatly decreased reproductivity in *japonica* rice.

We performed RNA-seq analysis in the young panicles of the following rice pairs: TP309-*Xa48* vs wild type TP309, and *CR-Xa48*/SKZ vs SKZ. The DEGs were enriched for glucose and amino acid metabolism pathways in *japonica* (Extended Data Fig. 8g, h), which may explain the defective grain development observed in in *japonica*. Taken together, these results suggest that the presence of *Xa48* decreases agronomic value with respect to reproduction and has been removed through artificial selection and mutation in *japonica*.

### A natural OsVOZ1 variant contributes to *Xa48*-induced yield penalty in *japonica* rice

The genetic mechanism underlying *Xa48* subspecies-specific selection was further explored by focusing on OsVOZ1, which contains a non-synonymous SNP, SNP^G/T^. This SNP resulted in an amino acid change from A to S (Fig. 4e). The allele frequency of *OsVOZ1^S^*was highly differentiated between *indica* and *japonica* rice and was examined during rice domestication and modern improvement. The allele frequency in wild rice was 5.0%, and increased to 38.6% during *indica* rice domestication, and continued to rise in modern *indica* cultivars (85.3%) (Fig. 4f, Supplementary Table 5). However, *OsVOZ1^S^* was rarely found in the *japonica* group (0.3%), which instead harbors the *OsVOZ1^A^* allele. In summary, the allele frequency changes of *Xa48* and *OsVOZ1^S^* suggested that the XA48-OsVOZ1 immune module may be favored and has been continuously co-selected during *indica* rice domestication and improvement. In contrast, *Xa48* and *OsVOZ1^S^* alleles in *japonica* rice have almost been eliminated possibly driven by purifying selection.

The SLC and Y2H assays demonstrated that both OsVOZ1^A^ and OsVOZ1^S^ interacted with XA48, indicating that this SNP does not impact the interaction of either OsVOZ1 variant with XA48 (Extended Data Fig. 9a, b). To determine the effect of *OsVOZ1* alleles on seed development, we randomly selected 70 *indica* accessions and divided them into four categories, namely OsVOZ1^S^ with or without *Xa48*, and OsVOZ1^A^ with or without *Xa48*. Seed setting rate of *indica* accessions were then assessed in a natural rice paddy. No significant difference was observed among these categories (Fig. 4g), suggesting that SNP^G/T^ has no effect on seed setting in *indica* rice. To further support this conclusion, the *indica* variety SKZ containing OsVOZ1^S^ was crossed with *indica* variety Kasalath containing OsVOZ1^A^ (Extended Data Fig. 9c); this *indica* × *indica* cross was not anticipated to impart significant changes in seed setting rate of the progeny, which generally occur in *indica* × *japonica* crosses. We identified four distinct groups of progeny populations by genotyping (*OsVOZ1^S^ Xa48*^-^, *OsVOZ1^A^ Xa48*^-^, *OsVOZ1^S^ Xa48*^+^, and *OsVOZ1^A^ Xa48*^+^); none showed difference in seed setting rate (Extended Data Fig. 9d). Moreover, we crossed SKZ with the *indica* varieties 9311 and TN1 (both containing *OsVOZ1^S^* and a mutated *Xa48*, Extended Data Fig. 9c) to obtain two OsVOZ1^S^-*Xa48*^-^ and OsVOZ1^S^-*Xa48*^+^ progenies and obtained similar results (Extended Data Fig. 9e). Additionally, when the *pXa48:Xa48* transgene was introduced into the *indica* varieties TN1 and ZS97 containing *OsVOZ1^S^* and *OsVOZ1^A^*, respectively, the resulting transgenic lines, TN1-*Xa48* and ZS97-*Xa48*, exhibited resistance to *Xoo* J18 but showed no difference in seed setting rate (Extended Data Fig. 9f-i). These results indicate that the XA48*-*mediated immunity module exhibits greater versatility in coping with different alleles of *OsVOZ1* in *indica* rice, while the XA48-OsVOZ1^A^ module specifically disturbs seed setting in *japonica*.

Our many attempts to edit the G/T SNP were unsuccessful, so we examined the function of this SNP in the seed setting rate of *japonica* using a transformation strategy. First, we generated lines overexpressing either allele (*OsVOZ1^A^*-OE or *OsVOZ1^S^*-OE) in NIPB using the maize *Ubiquitin* promoter, which showed no differences in seed setting relative to NIPB, but were significantly less resistant to *Xoo* (Fig. 4h, i), supporting the above notion that OsVOZ1 negatively regulates basal *Xoo* resistance (Fig. 3i). Next, we crossed the *OsVOZ1^A^*^/S^ overexpression lines with NIPB-*Xa48* to generate *OsVOZ1^A^*-OE/NIPB-*Xa48* and *OsVOZ1^S^*-OE/NIPB-*Xa48* plants, which displayed attenuated XA48-mediated resistance (Fig. 4j), in a manner consistent with results described above (Fig. 3g, h). The seed setting rate of *OsVOZ1^A^*-OE/NIPB-*Xa48* was lower than that in wild type NIPB. Interestingly, *OsVOZ1^S^*-OE/NIPB-*Xa48* restored the normal wild type seed setting rate (Fig. 4k, l). These results collectively demonstrate that the XA48-OsVOZ1^A^ module inhibits reproduction in *japonica*, and explain the asymmetric selection of the XA48-OsVOZ1 immune module between the two subspecies and the functional loss of *Xa48* in *japonica*.

### Breeding for BSR by integrating XA21-mediated PTI and XA48-mediated ETI

Our results demonstrated that *Xa48* confers a resistance spectrum that is complementary to that of *Xa21*, which originates from wild relative^11,12^. We proposed that the combination of *Xa48* and *Xa21*, which trigger ETI and PTI, respectively, may be a good approach that has practical implication for providing BSR in rice breeding programs.

This strategy was investigated by traditional cross breeding or transgenic approach (Extended Data Fig. 10a). The resulting *Xa21 Xa48* plants conferred BSR to 28 *Xoo* strains that were previously known to be avirulent on either *Xa21* or *Xa48* but not both (Fig. 5a, b, Extended Data Fig. 10b). Importantly, we established that stacking *Xa21* and *Xa48* conferred a high level of resistance to four *Xoo* strains (FJ-1∼4) that were virulent to both *Xa21* and *Xa48* (Fig. 5c); furthermore, this resistance was associated with increased expression of several pathogenesis-related (*PR*) genes during *Xoo* infection (Extended Data Fig. 10c). These results document that PTI and ETI pathways can be deployed together to enhance basal defense in rice. It is also noteworthy that the *Xa21 Xa48* rice lines retained a high level of *Xoo* resistance in the field trail after a flooding (Fig. 5d).

**Fig. 5.**
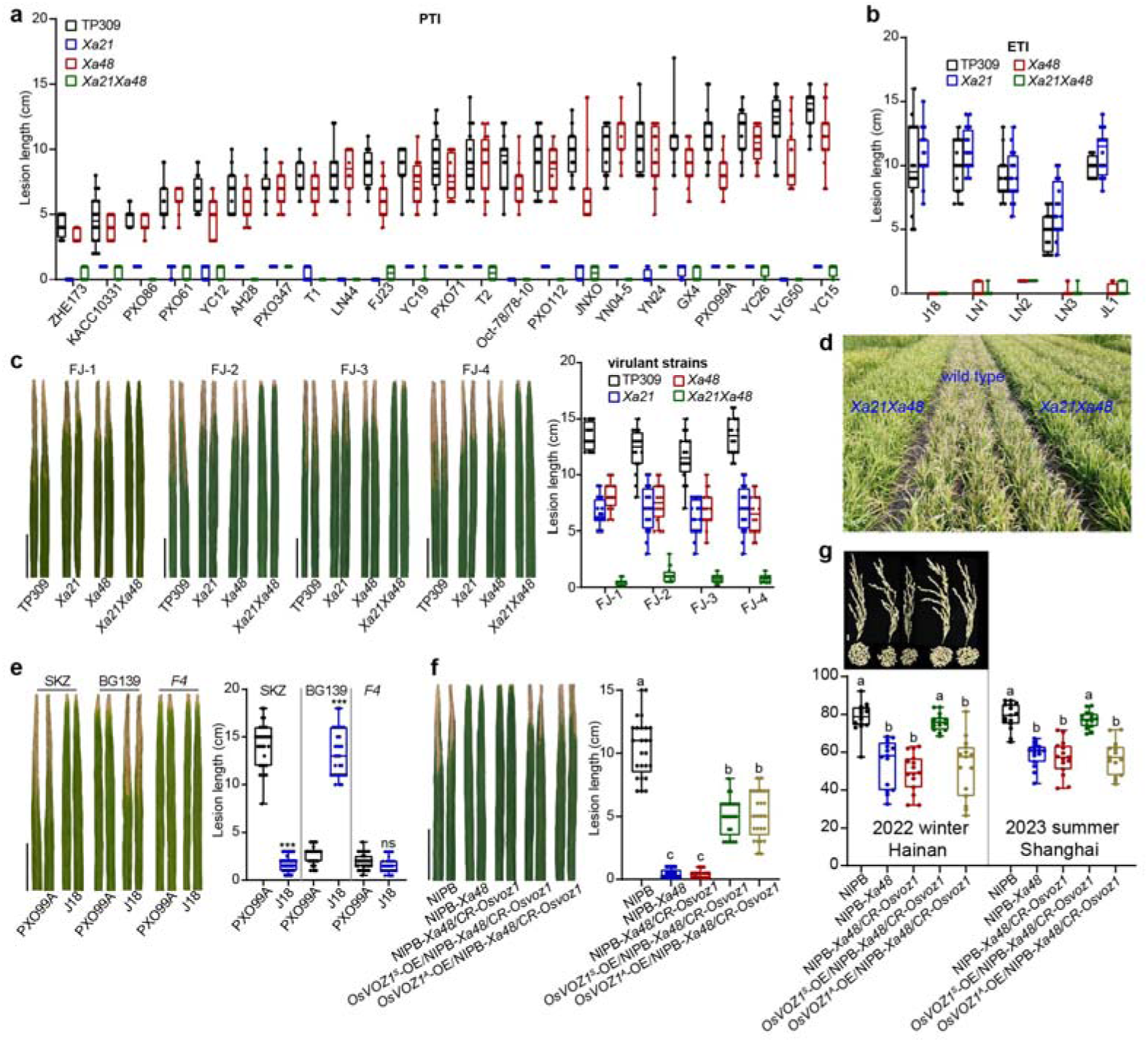
Broad-spectrum bacterial blight resistance is reconstructed by stacking. *Xa48*-mediated ETI and *Xa21*-mediated PTI in modern rice. **a**, **b**, Lesion lengths of TP309, *Xa21*(106), *Xa48*, and *Xa21Xa48* plants inoculated with 28 *Xoo* strains that are either virulent to *Xa48* or *Xa21*. Note that *Xa21Xa48* plants conferred broad-spectrum resistance in comparison to plants with the single *R* gene triggering PTI (**a**) and ETI (**b**) respectively. **c**, Improved resistance to strains FJ-1, FJ-2, FJ-3, and FJ-4 in *Xa21Xa48* plants. The strains were isolated from the infected leaves of rice plants grown in Fujian province (Southeast China), which bring a mutated XopG (Type V in Fig 2j) and are moderately virulent to respective *Xa21* and *Xa48*. **d**, *Xoo* nursery test of *Xa21Xa48* lines. Plants were kept under natural flooding after heavy rainstorms associated with a typhoon, leading natural *Xoo* infection in the Shanghai station. Disease symptom was recorded one and a half months after floodwaters. Note that *Xa21Xa48* rice conferred high filed resistance against *Xoo* infection. **e**, Representative leaves of SKZ and BG139 (*Xa21*) inoculated with PXO99A and J18. **f**, Lesion lengths of *japonica* regaining *Xa48* and OsVOZ1^S^ by crossing *CR-Osvoz1*/NIPB with NIPB-*Xa48*. **g**, De novo design of *japonica* rice restoring *Xa48* and OsVOZ1^S^. The *Xa48*-*OsVOZ1*^S^ *japonica* plants (*OsVOZ1^S^*-OE/*Xa48*/CR-*Osvoz1*) in NIPB background were grown in the paddy field for agronomic trait measurement with multiple-locations and seasons. Notably, the *Xa48*-*OsVOZ1*^S^ *japonica* plants restored normal seed setting. Data were shown as mean ± SD, *n* ≥ 15, asterisks represented statistical significance (\*\*\**P* < 0.001, two-tailed Student’s t-test). ns, not significant. Letters indicate significant differences (*P* < 0.05) determined by two-way analysis of variance (ANOVA) with Tukey’s test (**f**, **g**). Scale bars, 5 cm (**c**, **e**, **f**). Experiments were independently repeated three times with similar results (**a-c, e-g**).

Rice line BG139, an *indica* variety containing *Xa21*, was crossed with SKZ to generate *Xa21 Xa48* inbred *indica* lines; these contained intact *Xa21* and *Xa48* immune modules from wild rice and exhibited BSR to *Xoo* strains (Fig. 5e, Extended Data Fig. 10d). Because of the conventional approach for introducing the two genes, the utilization of these *Xa21 Xa48* lines in rice breeding programs is underway at several seed companies. In parallel, we *de novo* developed *japonica* lines expressing the *Xa48*-OsVOZ1^S^ immune module: we first knocked out *OsVOZ1^A^*in NIPB, the resulting *CR-Osvoz1* NIPB plants were then crossed with NIPB-*Xa48* to generate NIPB-*Xa48*/*CR-Osvoz1*. The later line was crossed again with *OsVOZ1^S^*-OE/NIPB to generate *OsVOZ1^S^*-OE/*Xa48* NIPB plants, which retained the XA48-OsVOZ1^S^ module and conferred resistance to *Xoo* J18 without a yield penalty (Fig. 5f, g, Extended Data Fig. 10e). It is important to mention that *Osvoz1* knockout plants were attenuated in growth and reproduction (Extended Data Fig. 10f), suggesting a critical function in plant growth and development. Collectively, our results provide a foundation for exploiting application of *R* genes from wild rice relatives with different immune modules and provides a breeding paradigm that high field disease resistance can be achieved through combining PTI and ETI in modern crop breeding.

## Discussion

Our study presents holistic approaches with *Xa48* for cultivating modern crops that exhibit both high disease resistance and optimal yield performance. First, leveraging *Xa48* and *Xa21* to reconstruct BSR to bacterial blight in rice, providing a success breeding example of how synchronized PTI and ETI indeed improves disease resistance in a real crop, as proposed^33–36^. This approach could also extend the shelf life of single *R* genes that have been defeated by virulent pathogen strains. Second, this study describes the use of genome editing, transgenes, and hybridization methods to created novel BLB-resistant, penalty-free *japonica* lines that recruit the XA48-OsVOZ1^S^ immune module. This breeding concept has the potential to be applied to other crop pathosystems, including rice for resistance to blast disease and wheat for resistance to stripe rust, utilizing reliable PTI and ETI modules.

XA48 and its associated transcription factor OsVOZ1 constitute an immune module that underwent differential artificial selection, leading to the loss of XA48 in *japonica* rice. This differential selection was driven by two main forces: one from the pathogen-host interaction, as presented with the XopG natural variation and distribution, and the other coming from artificial selection through selective breeding for high yield (Fig. 6a, b). Notably, the arms race of XA48 and XopG led to XopG enriched in strains from Northeast Asia where *japonica* rice is cultivated; In contrast, XopG had undergone massive loss-of-function events when *Xa48* is retained within the *indica* lineage, which has been widely cultivated in tropical and subtropical regions, encompassing the southern reaches of the Yangtze River and the Mekong River valley^37^, where typhoons and heavy rainstorms are common occurrences, as we observed in the natural *Xoo* nursery (Fig. 6a). This specific agroecological scenario, coupled with high pathogen pressure, is likely a contributing factor to the retention of *Xa48* in *indica* rice.

**Fig. 6.**
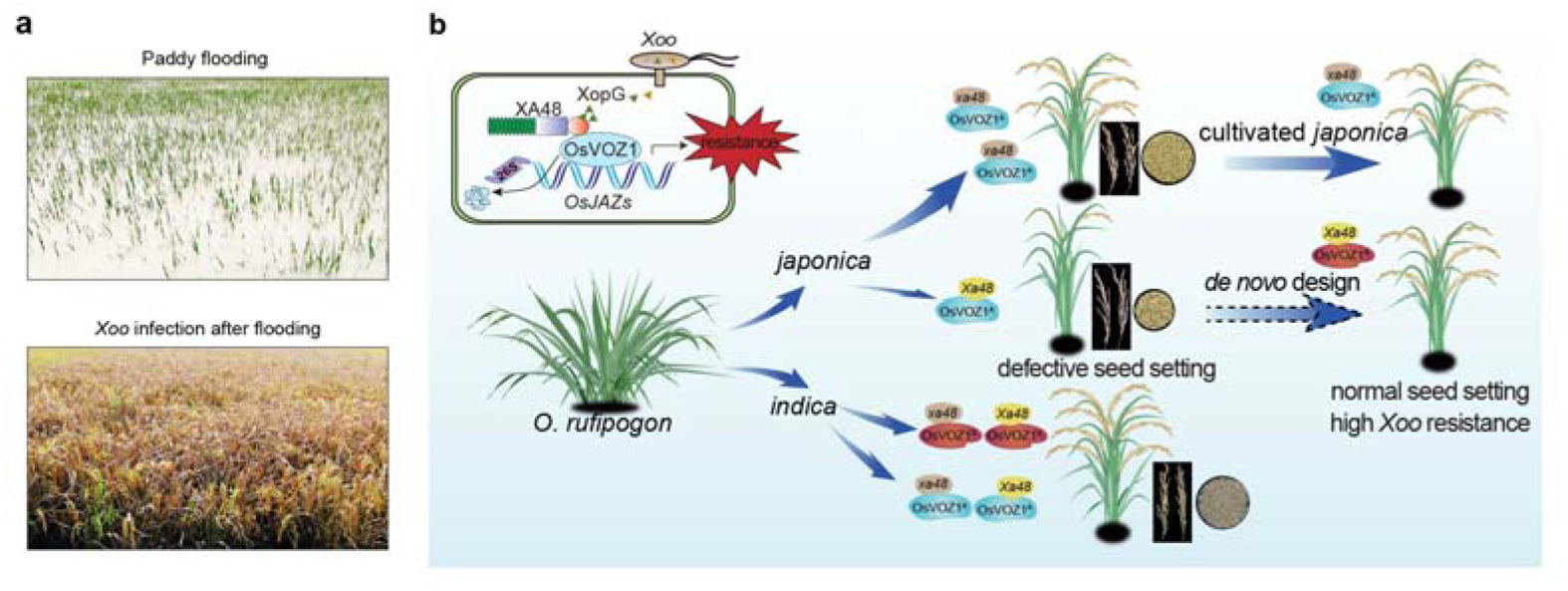
**Working model of subspecies-specific selection of the XA48-OsVOZ1 module and design in *japonica***. **a**, Rice leaf bacterial blight spread after floodwaters associated the typhoon in the Shaoxing experimental station (Southeast China) in the 2021 summer, showing paddy flooding (top, July) and bacterial blight symptom in the field (bottom, September). **b**, A proposal model for subspecies-specific selection of the XA48-OsVOZ1 immune module between *indica* and *japonica*. XA48 perceives the cognate type III bacterial effector XopG to activate immunity and promote degradation of the immunosuppressive transcription factor OsVOZ1, which activates *OsJAZs*, the suppressors of immunity. *OsVOZ1* evolved two alleles, *OsVOZ1^A^*and *OsVOZ1^S^*, *japonica* rice only inherits *OsVOZ1^A^* while *indica* rice keeps both. The XA48-OsVOZ1^A^ combination poses a negative effect on reproductivity in *japonica*, leading to *Xa48* functional loss in *japonica*. *Xa48* is kept and does not affect reproduction in *indica* with either OsVOZ1^A^ or OsVOZ1^S^ allele, which is historically cultivated in Southeast Asia where *Xoo* frequently outbreaks due to typhoons and floodwaters.

The reintroduction of XA48 in *japonica* lines resulted in reduced seed production due to combination of *Xa48* and the *japonica* allele *OsVOZ1^A^*, while the *Xa48* immune module is much more versatile in coping with different alleles of *OsVOZ1 ^A/S^* in *indica* (Fig. 6b). The immune system incompatibility is reminiscent of hybrid necrosis or weakness seen in plants, where genetic incompatibility triggers autoimmunity in hybrids. This autoimmunity primarily arises from a conflict involving NLRs, which are crucial for achieving a compatible immune response and better yield performance in hybrids but become problematic when their interactions lead to incompatibility^38–41^. Therefore, our study documents that the incompatible immune system has led to the artificial selection for an ETI network that shapes both disease resistance and reproduction in a crop.

## Methods

### Plant materials and growth conditions

The *japonica* model varieties NIPB, TP309 and transgenic TP309 line 106 expressing *Xa21*, the *indica* varieties, SKZ, Kasalath, ZS97, 9311 and TN1, and 1,945 available rice accessions used in the 3K genome project^27^, were used and grown in the experimental paddy fields at Shanghai for the summer season and Hainan Island for the winter season under natural field conditions. *Nicotiana benthamiana* grown at a growth chamber under long-day conditions (16-h day/8-h night, 22°C) for 3-4 weeks, was used for transient expression experiments, protein-protein interaction, and cell death assays.

### Pathogens

*Xoo* strains (listed in Supplementary Table 6) from Asia and elsewhere were stored and used in this study. *Xtt* Km9, *Pst* DC3000, *Psa* M228, *E. coli* O157 and *Sea* SA20100345 were also used in this study.

### Map-based cloning and screening of rice BAC library

SKZ was crossed to NIPB to generate an F_2_ mapping population and inoculated with *Xoo* strain J18. *Xa48* was delimited to a 65-kb genomic region between InDel33 and InDel31 on chromosome 3. The genomic DNA of SKZ was partially digested with *Hin*dIII and inserted into the BAC cloning vector plndigoBAC-5 for library construction. A BAC clone was selected that spanned the *Xa48* locus. Sequence analysis of the BAC clone revealed four ORFs within the 65-kb region, which showed several insertion, deletion and SNPs between NIPB and SKZ.

### GWAS analysis of *Xoo* resistance

The genotyping matrix for GWAS was obtained from the 3K Rice Genomes Project^27^ and other rice genome projects^37,42^. The panel includes 4,591 rice accessions with 5.23 million SNPs. 1,945 accessions were phenotyped for J18 resistance, a total of 3,605,578 SNPs with minor allele frequency above 0.05 were maintained for the GWAS study. GWAS was performed by GEMMA with the linear mixed model^43^. Kinship matrix was incorporated in the GWAS models to control false-positives. The effective number of SNPs was calculated as 786,858 by GEC software^44^, and therefore by applying genome-wide type I error rate at α□=□0.05, we determined the significance threshold as 6.35E-8 (0.05/786,858). The GWAS plot was visualized using the CMplot R package^45^.

### Allele frequency inference during rice domestication and improvement

A total of 2,451 Asian rice accessions were used for inference of the allele frequency changes of Os*VOZ1* and *Xa48*. These accessions include 169 wild rice (88 and 81 genetically close to *indica* and *japonica*, respectively), 1,158 temperate *japonica*, 135 tropical *japonica*, 22 *basmati*, 79 *aus* and 888 *indica* rice^29,30^. Detailed information is listed in the Supplementary Table 3. Landraces and cultivar accessions for *indica* and temperate *japonica* were classified based on Wei et al. (2021). Variants in the Os*VOZ1 and Xa48 genes* were called by GATK v3.7^46^. The allele frequency was calculated for variants in *Xa48* and Os*VOZ1* for all rice groups.

### Plasmid construction and rice transformation

For CRISPR/Cas9-mediated knockout mutation in rice, 20-bp gene-specific guide RNA sequences were constructed targeting the candidates *C1*, *C2*, *C3*, *C4* of the mapping region and *OsVOZs* were PCR-based cloned and subcloned into pYLCRISPR/Cas9-MH vector^47^. To generate the constructs for *OsVOZ1* and *OsVOZ2* fusions driven by the maize *Ubiquitin 1* promoter (Ubi), coding regions (CDS) were inserted into PUN1301-pUBI-GFP or PUN1301-pUBI-FLAG vectors. For complementation, DNA fragments containing the whole *Xa48* promoter and coding region were inserted into binary vector pCAMBIA1300 to generate plasmid *pXa48::Xa48*.

All constructs were introduced into *Agrobacterium* strain EHA105, and then transformed into different rice backgrounds to generate more than 15 independent lines; an exception was the CRISPR/Cas9 knockout mutants that usually generated more than 100 transformants for mutation screening. All transgenic plants were selected by PCR-based sequencing, gene expression assays or western blot. For transient expression in *N. benthamiana*, the constructs were introduced into *Agrobacterium* strain GV3101. The primer sequences used for cloning are listed in Supplementary Table 7.

The following transgenic lines/mutants were developed in this study: *CR-C1* (*CR-Xa48*/SKZ), *CR-C2*, *CR-C3*, *CR-C4*, NIPB-*Xa48*, TP309-*Xa48*, Kasalath-*Xa48*, TN1-*Xa48*, ZS97-*Xa48*, *OsVOZ1*-OE/SKZ, *OsVOZ2*-OE/SKZ, *OsVOZ1*-OE/NIPB-*Xa48*, *OsVOZ2*-OE/NIPB-*Xa48*, *OsVOZ1^A^*-OE, *OsVOZ1^S^*-OE, *OsVOZ1^A^*-OE/NIPB-*Xa48*, *OsVOZ1^S^*-OE/NIPB-*Xa48*, *xopG*-OE, *XA48*-OE, *CR-Osvoz1/*NIPB, *CR-Osvoz2/*NIPB, NIPB-*Xa48/CR-Osvoz1*, *OsVOZ1^S^*-OE/NIPB-*Xa48/CR-Osvoz1*, *OsVOZ1^A^*-OE/NIPB-*Xa48/CR-Osvoz1* and *Xa21*/*Xa48*. All transformants were grown in experimental paddy fields located at Shanghai and Hainan Island.

### Pathogen inoculation

The *Xoo* strains used in this study included strains from Northeast Asia, South Asia and other regions, and *Xoo* transformants expressing *xopG* homologs from other pathogenic bacteria (listed in Supplementary Table 6). For inoculation experiments, *Xoo* strains grown on PSA medium (10 g/L tryptone, 10 g/L sucrose, 1g/L glutamate, 15 mg/L cephalexin, pH 7.0) at 28□ for 3 days were suspended in sterile water to OD_600_ = 1.0. Two-month-old plants were inoculated by leaf-clipping and syringe infiltration methods. Lesion length was measured at 14 days post inoculation (dpi). *Xoo* growth curve were generated by suspending inoculated leaf tissue (20 cm) in 10 ml sterile water to collect bacteria; suspensions were diluted to count colony-forming units. Inoculation experiments were repeated independently at least three times.

### Rice *Xoo* nursery test

To investigate *Xoo* infection and potential epidemic after flooding, rice was grown at the Shaoxing (Zhejiang province) and Shanghai experimental station in June, 2021. Plants were maintained for 5 d under natural flooding due to August rainstorms. Plants were maintained without bactericide application to allow *Xoo* to infect naturally. Disease symptoms were recorded 45 d after flooding.

### Tn*5* insertion mutagenesis of *Xoo*

Competent *Xoo* LN2 cells were mixed with 1 μL of the EZ-Tn*5* <R6Kγori/KAN-2>Tnp Transposome and subjected to electroporation as recommended by the manufacturer (Epicentre, Madison, USA). The transformed *Xoo* cells were transferred to 1.5 mL Eppendorf tubes, incubated at 180 rpm at 28°C for 1.5 h, and then cultured on PSA containing the kanamycin (20 μg/mL) at 28°C for 3 d. Over 20,000 Km-resistance colonies were individually numbered and then transferred to new PSA plates with Km for further investigation.

### Plasmid rescue and XopG identification

*Xoo* genomic DNA was extracted by the CTAB method and digested. The digested DNA was separated in 1.3% agarose gels at 80V for 16-20 h and then transferred to Immobilon-Ny+ Transfer Membranes (Merck, USA). The DNA was labeled, hybridized, and detected using the DIG-High Prime DNA Labeling and Detection Starter Kit. Plasmid rescue was performed with the EZ-Tn*5*^TM^ <R6Kγori/KAN-2>Tnp Transposome^TM^ kit following the manufacturer’s instructions.

For XopG identification, the genomic DNA was isolated from two Tn*5* mutants of *Xoo* strain LN2, which were compatible with NIPB-*Xa48*. The genomic DNA was extracted, digested with *Pst*I enzyme, and hybridized in Southern blots where Tn*5* was used as a probe. Detection was conducted with the Dig-Labeling Kit (Roche) as recommended by the manufacturer. The *Pst*I-digested gDNA fragments were also self-ligated using T_4_ ligase and transformed into competent cells of *E. coli* EC100D (Lucigen, USA). The plasmids of single clone were isolated and fragments flanking the Tn*5* insertion site were sequenced using the primers included in the Tnp Transposome kit. The sequenced fragments were analyzed using BLAST programs available at the National Center for Biotechnology Information (NCBI).

### Expression of *xopG* homologues from other bacterial pathogens

Both *xopG* and *xopG* homologues were PCR-amplified (see primers in Supplementary Table 7) and ligated into pHM1 vector at the *Sal*I-*Hin*dIII site. Competent cells of *Xoo* PXO99A were electroporated with 1 μL purified plasmid DNA and transferred to NA containing spectinomycin. The transformed *Xoo* PXO99A strains were verified by immunoblotting using FLAG as the antibody.

### Yeast two-hybrid screening and protein interaction analysis

Y2H screening was conducted to identify XA48-interacting proteins with a rice pathogen-induced cDNA library as previously described^22^. For the Y2H assay, the coding sequences of target genes, including *xopG*, were amplified with gene specific primers and cloned into yeast expression vectors pDEST22 (AD)/32 (BD) (Invitrogen). Constructs were co-transformed into yeast strain AH109, grown on selective medium (lacking Trp and Leu or Trp, Leu and His) containing 3-aminotriazole (Sigma-Aldrich, A8056) for 3 d at 30□.

### Split luciferase complementation assays

For SLC assays, coding sequences of the target genes were cloned into pCAMBIA-35S-nLuc or pCAMBIA-35S-cLuc, which were transformed into *Agrobacterium* strain GV3101. *Agrobacterium* cells were collected and resuspended in infiltration buffer (10 mM MgCl_2_, 10 mM MES, 150 mM acetosyringone, pH 5.6), and incubated for 2-3h at 30°C before infiltration into *N. benthamiana* leaves. Two days after transformation, luciferase activity was measured as recommended by the manufacturer (Promega), images were captured using the Tanon-5200 Chemiluminescent imaging system (Tanon).

### Bimolecular fluorescence complementation assay

For BiFC assays, XopG and XA48 CC were fused to the N-terminal fragment of YFP (nYFP) and the C-terminal fragment of YFP (cYFP), which were transformed into *Agrobacterium* strain GV3101. *Agrobacterium* cells were collected, resuspended in infiltration buffer, and incubated for 2-3 h at 30°C before infiltration into *N. benthamiana* leaves. Fluorescence images were harvested at 48 post-infiltration (hpi) using a Leica TCS SP5 II confocal microscope.

### Protein co-immunoprecipitation

Protein co-IP assays were performed to verify protein-protein interactions *in planta*. *Xa48-CC* was cloned to pCAMBIA1300-35S-eGFP(C)-rbcsE9 and *OsVOZ1*, *OsVOZ2* and *xopG* were cloned to pCAMBIA1300-35S-FLAG(C)-rbcsE9. Protein extracts were prepared from *N. benthamiana* or transgenic rice leaves in IP buffer [50 mM Tris-HCl, pH 7.5, 150 mM NaCl, 1 mM EDTA, 10% glycerol, 1% Triton X-100, 1mM PMSF and protease inhibitor cocktail (11836153001, Sigma)]. Supernatants were incubated with anti-GFP beads for 2 h at 4°C and then washed four times with washing buffer (50 mM Tris-HCl, pH 7.5, 150 mM NaCl, 1 mM EDTA, 10% glycerol, 1 mM PMSF and protease inhibitor cocktail). The bound proteins were released from the beads by boiling for 5 min in SDS loading buffer. The reagents used included the antibodies, FLAG (Sigma, F1804), GFP (Abcam, ab290), goat anti-rabbit IgG secondary antibody (Thermo Fisher, 31460) and goat anti-mouse IgG secondary antibody (CWBIO, CW0102).

### Subcellular localization

For protein subcellular localization assay, coding sequences of the genes were cloned into PA7-35S-YFP or PA7-35S-mCherry, which were transformed into rice protoplasts prepared from leaf sheaths of 10-day-old seedlings. NLS (Nuclear Localization Signal) was used as a nuclear marker and OsRAC1 as a plasma membrane marker^22^. After incubation in the dark for 16-20□h at 26°C, fluorescence was observed with a confocal microscope (Zeiss LSM 880).

### Cell death assay in *N. benthamiana*

The *N. benthamiana* cell death assay was used to determine the function of XA48-XopG interaction and XA48 activation. XopG-GFP and XA48-GFP constructs were transformed into *Agrobacterium* GV3101 and expressed in *N. benthamiana*. The expression of each protein was confirmed by immunoblotting using anti-GFP and cell death on infiltrated leaves was photographed at 36-48 hpi.

### Cell-free degradation assay

Total proteins were extracted from *N. benthamiana* leaves or transgenic rice leaves with the extraction buffer (50mM Tris-HCL, pH 7.5, 100 mM NaCl, 10 mM MgCl_2_, 5 mM dithiothreitol, 5 mM adenosine 5‘-triphosphate, and 1x protease inhibitor cocktail). The extracts were clarified by two sequential centrifugations at 12,000 g for 5 min. Samples were incubated at 25 °C at the indicated time points. The reaction was terminated by boiling the reaction in SDS sample buffer and then analyzed by western blotting. All protein immunodetection experiments were performed independently three times with similar results.

### GUS reporter assay

For the β-glucuronidase (GUS) reporter gene constructs, the 3-kb promoter region of *Xa48* was inserted upstream of the GUS coding sequence in expression vector pCAMBIA1300-GUS and then introduced into *Agrobacterium* strain EHA105 to generate GUS reporter transgenic plants.

For GUS staining, plant tissues were incubation-stained in buffer (50 mM NaPO_4_, pH 7.0, 5 mM K_3_Fe(CN)_6_, 5 mM K_4_Fe_6_, 0.1% Triton X-100, and 1 mM X-Gluc) overnight at 37°C, and then dehydrated with a graded ethanol series.

### RNA preparation and RNA-seq analysis

Total RNAs were extracted using TRIzol reagent according to the manufacturer’s instructions (Invitrogen). For quantification of gene expression, 1□μg RNA was reverse-transcribed into cDNA using oligo (dT) primers and SuperScript III. qRT-PCR was performed using SYBR Premix Ex Taq (TaKaRa) and gene-specific primers were listed in Supplementary Table 7, rice *ACTIN* served as an internal control to normalize expression levels. The 2^-△△CT^ method was used to calculate the relative expression levels with three biological repeats.

For RNA-seq analysis, RNA samples were prepared from *Xoo*-infected rice leaves at 24 hpi and from panicles of different rice genotypes at the booting stage. RNA-seq was performed by Shanghai Biotechnology Corporation. Three biological replicates were used for each genotype/treatment. The entire RNA-seq dataset was deposited in the NCBI Gene Sequence Read Archive (SRA) under accession number: PRJNA1028151.

### CUT& Tag assay

Cleavage Under Targets and Tagmentation (CUT&Tag) assays were conducted using the Hyperactive Universal CUT&Tag Assay Kit (Vazyme, Cat# TD903). Briefly, two-week-old seedling leaves of OsVOZ1-OE/NIPB and OsVOZ1-OE/SKZ were cross-linked with 1% formaldehyde and stopped by glycine solution. The tissue was ground into fine powder and nuclei were prepared in NE1 buffer (20□mM HEPES, pH 7.5, 10□mM NaCl, 0.5□mM spermidine, 0.1% Triton X-100, 20% glycerol) for 10□min on ice. Samples were resuspended in NE2 buffer (20□mM HEPES, pH 7.5, 150□mM NaCl, 0.5□mM spermidine, Protease Inhibitor) and centrifuged at 3000 g for 5□min. Subsequently, the anti-GFP antibody (Abcam, Cat#ab290) was added and incubated overnight at 4°C, accompanied by gentle rotation. Secondary antibody was added and incubated for 1□h at room temperature. DNA library was constructed using TruePrep Index Kit V2 for Illumina (Vazyme, Cat# TD202) with proper primers.

### Electrophoretic mobility shift assay

OsVOZ1-MBP recombinant protein was developed and purified in *E. coli*. DNA fragments were end-labeled with Cy5, which was incubated with 0.2 ug protein in 20 ul binding buffer (100 mM Tris-HCl, pH 7.5, 250 mM KCl, 25 mM MgCl_2_, 10 mM EDTA, 10 mM DTT and 10% glycerol) at room temperature for 30 min. 50-200-fold non-labeled competitor DNA was added for competition assays. The reaction mixture was electrophoresed on a native polyacrylamide gel, and then was imaged by autoradiography (Fujifilm FLA 9000 plus DAGE) according to the manufacturer.

### Dual-luciferase transcriptional activity assay

A dual-Luc reporter system in *N. benthamiana* leaves was performed following a previously reported protocol^22^. Briefly, pCAMBIA1300-35s-GFP and pCAMBIA1300-35s-OsVOZ1-GFP serve as effectors. 3 kb promoter region of *OsJAZs* were cloned into pGreenII-0800-Luc vector to generate Pro*OsJAZ*::Luc reporter. The effector and reporter plasmids were introduced into *Agrobacterium* strain GV3101 and co-expressed in *N. benthamiana* leaves. The LUC and REN luciferase activities were measured using Dual-Luciferase Reporter Assay Kit (Promega) following the manufacturer’s instructions.

### Phylogenetic analysis

For phylogenetic analysis of XopG, we utilized the amino acid sequence of XopG as a query to identify homolog proteins in other Gram-negative bacteria via the Domain Enhanced Lookup Time Accelerated BLAST (BLASTp) program available at NCBI. Subsequently, XopG and its identified homologs were employed to construct a phylogenetic tree using the Neighbor-Joining method, with 1,000 bootstrap replicates for statistical support, in MEGA11 software.

### Breeding approaches for stacking *Xa***21** and *Xa***48**

We used two approaches to stacking *Xa21* and *Xa48*. First, transgenic TP309-*Xa21*(106) was crossed to TP309-*Xa48*, and the progeny were self-crossed continually to generate homozygous polymeric *Xa21Xa48* plants (F_5_). In a second approach, the traditional *indica* variety BG139 (containing *Xa21*) was crossed to SKZ (containing *Xa48*), and the progeny self-crossed to generate homozygous breeding material *Xa21 Xa48* (F_4_).

### De novo development of *japonica* rice expressing *Xa*48 and *OsVOZ*1*^S^*

To re-introduce *Xa48* and *OsVOZ1^S^* into *japonica* rice, we first generated *OsVOZ1-*KO lines in NIPB plants (*CR-Osvoz1*). We then crossed *CR-Osvoz1* to NIPB-*Xa48* to develop NIPB-*Xa48/CR-Osvoz1*, and crossed to *OsVOZ1^S^*-OE to develop *OsVOZ1^S^*-OE/NIPB-*Xa48/CR-Osvoz1* plants. The *Xa48-OsVOZ1^S^ japonica* plants (*OsVOZ1^S^*-OE/NIPB-*Xa48/CR-Osvoz1 NIPB*) were grown in the paddy field for measurement of agronomic traits.

### Field trails and agronomic trait measurements

One-month-old seedlings were transplanted into paddy fields located at the Shanghai and Hainan Experiment Stations during 2021∼2023 for multiple-season and-location field test. At least 12 independent lines of each genotype were grown over multiple seasons at both locations. Upon maturity, agronomic traits including plant height, tiller number, seed setting rate, grain number per panicle, grain weight per plant, and 1,000-grain weight were analyzed. Average values of 15 plants or 12 representative progeny lines of each genotype (T3) were measured and compared.

## Statistical analysis

Quantification analysis on lesion length, bacterial growth, plant growth, seed setting rate, grain productivity and other measurements were conducted in GraphPad Prism. All values were presented as mean ± SD and the number (n) of samples or replicates were indicated in the corresponding figure legends. Significance of difference was analyzed using Student’s t test for pairwise comparisons and one-way or two-way ANOVA followed by Tukey’s test for multiple groups’ comparison. Detailed information about quantifications and statistical analysis values is provided in figure legends, or within specific sections of Methods. No methods were used for sample randomization or sample size estimation and no data were excluded from analysis.

## Data availability

The RNA-seq data generated in this study have been deposited in the SRA database under accession PRJNA1028151. The full genomic sequence of the *Xa48* can be found in GenBank 2754159. All data are available in the main text or the supplementary materials. Source data are provided with this paper.

## Acknowledgements

We would like to thank Dr. Qifa Zhang for helpful discussion, Dr. Pamela Ronald for providing the *Xa21* transgenic seeds, Ms. Xin Wang for rice transformation, Ziyao Lei for field inoculation, and Mr. Jianyao Shou for helping with *Xoo* nursery. This work was supported by Biological Breeding-National Science and Technology Major Projects (2023ZD04070 to Y.D.), the National Natural Science Foundation of China (32088102, 31930090 to Z.H.; 32361143515, 31830072 to G.C.; U20A2021 to Y.D.; 32402392 to H.L), the Chinese Academy of Sciences (XDB27040201 to Z.H. and XDA24010304 to Y.D.), Shanghai Science and Technology Development Funds (24YF2751900 to H.L), Shanghai Agricultural Science and Technology Innovation Program (Grant No. K2024-02-08-00-12-F00050 to H.L), and the National Key Research and Development Program of China (2022YFE0198100 to Y.D.).

## Author contributions

H.L., F.C., Y.D., G.C., and Z.H. conceived and designed the experiments. H.L., F.C., B.Y., G-Y.C., M.Y., and Y.C. performed experiments, and Y.W. and G-Y.C. performed Tn5 mutagenesis. J.Q., X.H. and B.H. performed population genomics and artificial selection analysis. K.C., X.G., S.L., J.L., and J-S.J. assisted with pathogen inoculation and field trials. B.M. assisted in rice transformation experiments. R.L. J.W. and J.X. grew and provided wild rice and rice germplasm. Z.H., Y.D. and G.C. supervised the project. Z.H. and Y.D. provided theoretical contributions to the project. H.L., F.C., Y.D., G.C. and Z.H. analyzed the data and wrote the paper. All authors discussed and commented on the manuscript.

---

**Extended Data Fig. 1.**
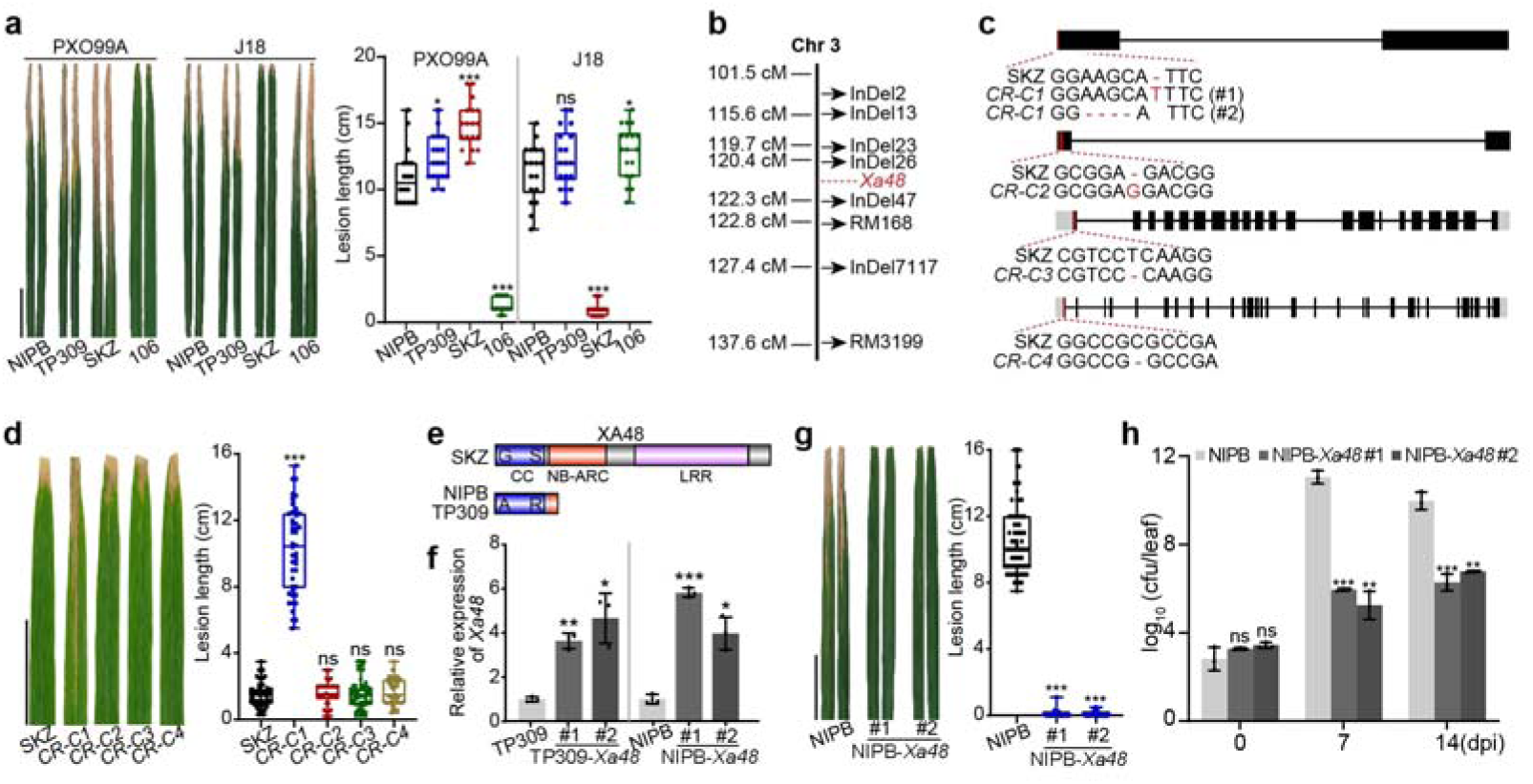
Map-based cloning identifies *Xa48*. **a**, SKZ displayed strong resistance to J18 but high susceptible to PXO99A. Disease symptoms of rice variety SKZ, NIPB, TP309, and 106 at 14 dpi with *Xoo* strains J18 and PXO99A. Line 106 is transgenic TP309 expressing *Xa21*. **b**, *Xa48* was initially mapped to chromosome 3. **c**, **d**, Independent CRISPR/Cas9 knockout lines (*CR-C1*, *CR-C2*, *CR-C3*, *CR-C4*) of SKZ were generated (**c**), and representative lines’ lesion length of knockout mutants were measured (**d**). Note that only *CR-C1* became susceptible to J18. **e**, Premature termination of XA48 translation in susceptible varieties caused by a single-base deletion and SNP. Schematic of the domain structure of XA48 is indicated. CC, coiled coil; NBS, nucleotide binding sequence; LRR, leucine-rich repeat. **f**, Transcriptional level of *XA48* was detected in TP309-*Xa48* (**f**) and NIPB-*Xa48* (**g**) complement plants. **g**, **h**, NIPB-*Xa48* transgenic plants are highly resistant to J18 as SKZ. The lesion length (**g**) and bacterial populations (**h**) were measured. Data were shown as mean ± SD, *n* ≥ 30 (**a, d, g**) and *n* = 3 (**f, h**), asterisks represented statistical significance (\**P* < 0.05, \*\**P* < 0.01, \*\*\**P* < 0.001, two-tailed Student’s t-test). ns, not significant. Scale bars, 5 cm (**a, d, g**). Experiments were independently repeated three times with similar results (**a, d, g**).

**Extended Data Fig. 2.**
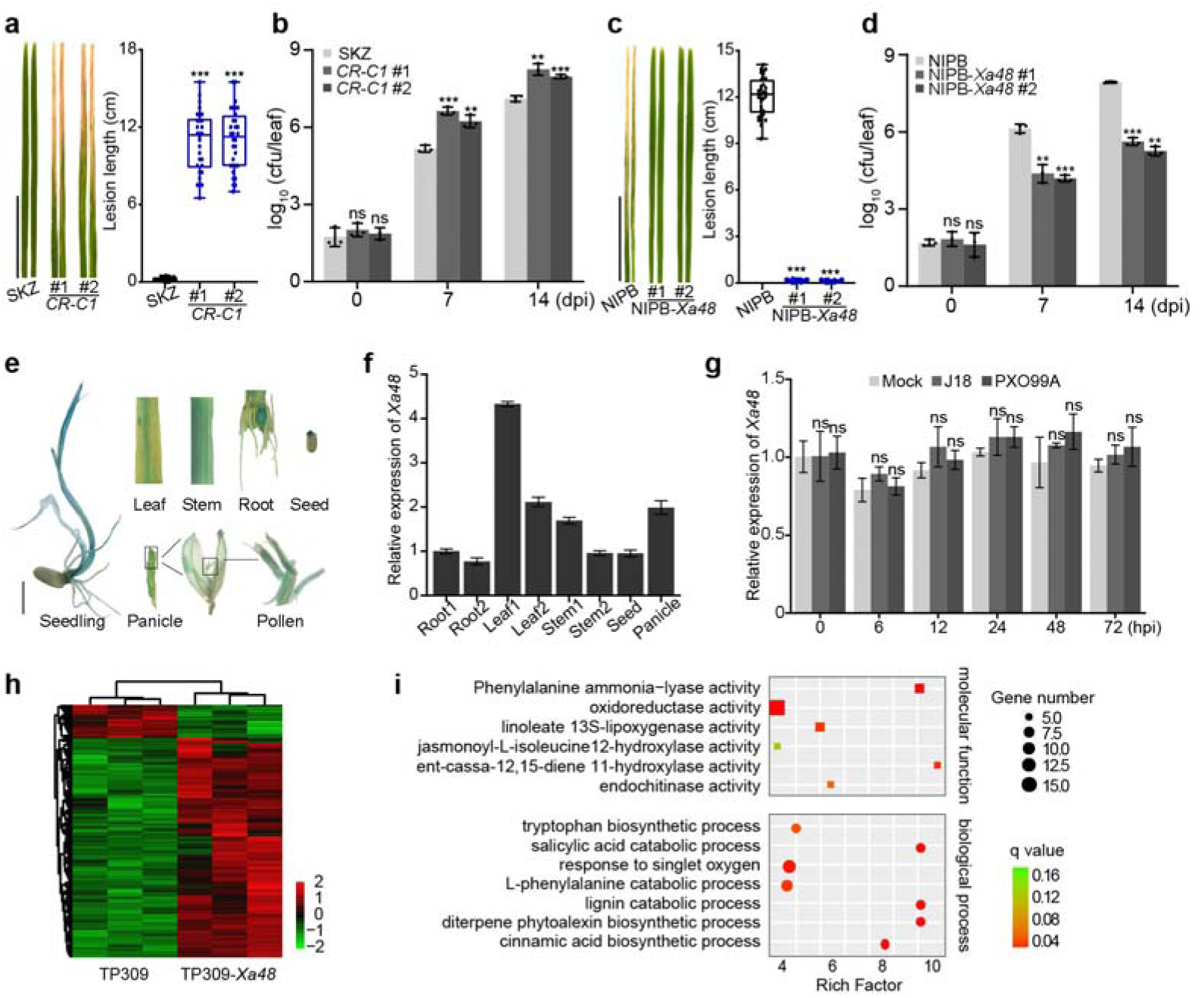
***Xa48* confers lifetime *Xoo* resistance. a-d**, *Xa48* confers *Xoo* resistance in SKZ (*indica*) (**a**, **b**) and NIPB-*Xa48* (*japonica*) (**c**, **d**) seedlings. Lesion lengths and bacterial populations of representative *CR-Xa48*/SKZ mutants and NIPB-*Xa48* were measured in seedlings inoculated with J18. **e**, GUS staining of *pXa48::GUS* reporter transgenic plants revealed *Xa48* expression in leaf, stem, root, seed, and young panicle. The boxed sections were magnified for close views of GUS staining in the spikelet and anther. Scale bars, 1 cm. **f**, Expression patterns of *Xa48*. Expression of *Xa48* in different tissues was detected by qRT-PCR and normalized to expression in the root 1. Root1 was from 5-day-old seedling, leaf1, stem1 were from 25-day-old seedling, and root2, leaf2, stem2 were from 80-day-old plants. **g**, *Xa48* was not induced by *Xo*o infection as revealed by qPCR. Four-week-old rice were inoculated with J18 and PXO99A, and leaf samples were collected during a time course of 0-72 hpi for RNA preparation. **h**, **i**, Hierarchical clustering and Gene Ontology (GO) analysis of differentially expressed genes in TP309 and TP309-*Xa48* leaves inoculated with J18. Note that *Xa48* enhanced gene induction in salicylic acid, ethylene, and diterpene phytoalexin biosynthesis processes, which play important roles in rice immunity. Data were shown as mean ± SD, *n* ≥ 30 (**a, c**) and *n* = 3 (**b, d, f, g**), asterisks represented statistical significance (\*\**P* < 0.01, \*\*\**P* < 0.001, two-tailed Student’s t-test). ns, not significant. Experiments were independently repeated three times with similar results (**a, c**).

**Extended Data Fig. 3.**
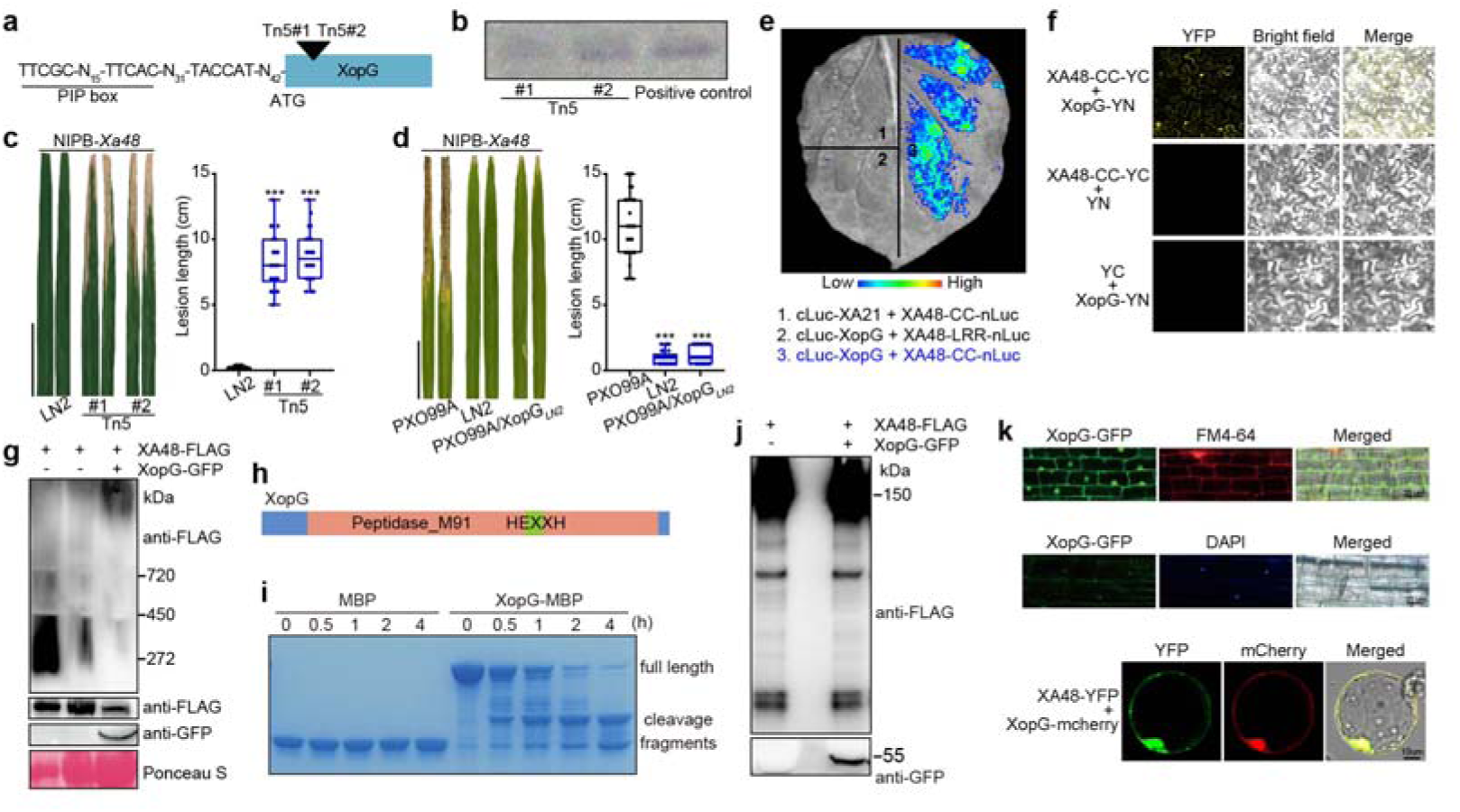
Identification and functional analysis of XopG through Tn*5* mutagenesis. **a**, **b**, Generation of *avrXa48*/*xopG* insertion mutants. Tn*5* transposon-insertion shows two independent insert mutants (#1 and #2) in the same site (**a**), which cause loss-of-function of XopG. Southern blot confirms single insertion of Tn*5* in the two LN2 mutants (**b**). The non-TAL promoter of *XopG* is indicated with the featured sequence. **c**, Tn*5* transposon-inserted mutants of strain LN2 lost avirulence to NIPB-*Xa48*. The lesion length was measured at 14 dpi. **d**, The introduction of XopG*_LN2_* changed PXO99A into avirulence to NIPB-*Xa48*. **e**, XopG directly interacts with the XA48 CC domain by SLC in *N. benthamiana*. XA21 and XA48-LRR domains, which do not interact with XopG were used as negative controls. **f**, XopG associates with the XA48 CC mainly in the cell periphery and nucleus as revealed by BiFC in *N. benthamiana* leaf cells. XopG and XA48-CC were fused to the N-terminal fragment of YFP (nYFP) and the C-terminal fragment of YFP (cYFP). Scale bars, 50 μm. **g**, XopG likely induces oligomerization of XA48 in rice cells as revealed by BN-PAGE assay. Total protein was detected by immunoblotting with anti-FLAG and-GFP antibodies. **h**, Schematic of the domain structure of XopG is indicated. Peptidase_M91 contains an HEXXH motif, characteristic of zinc metallopeptidases. **i**, **j**, XopG displayed self-cleavage activity (**i**) but did not cleavage XA48 (**j**). Immunoblots of XopG-MBP and MBP acts as control. XA48-FLAG with XopG-GFP detected by immunoblotting with anti-FLAG/GFP antibodies. **k**, Representative localization images of XopG-GFP transgenic rice, showing main localization to the PM and nucleus. Co-localization of XA48 with XopG was detected in the PM and nucleus transiently expressed in rice protoplasts. FM4-64 and DAPI staining indicate the PM and nucleus. Data were shown as mean ± SD, *n* ≥ 22 (**c, d**), scale bars, 5 cm, asterisks represented statistical significance (\*\*\**P* < 0.001, two-tailed Student’s t-test). Experiments were independently repeated three times with similar results (**c, d, e, j**).

**Extended Data Fig. 4.**
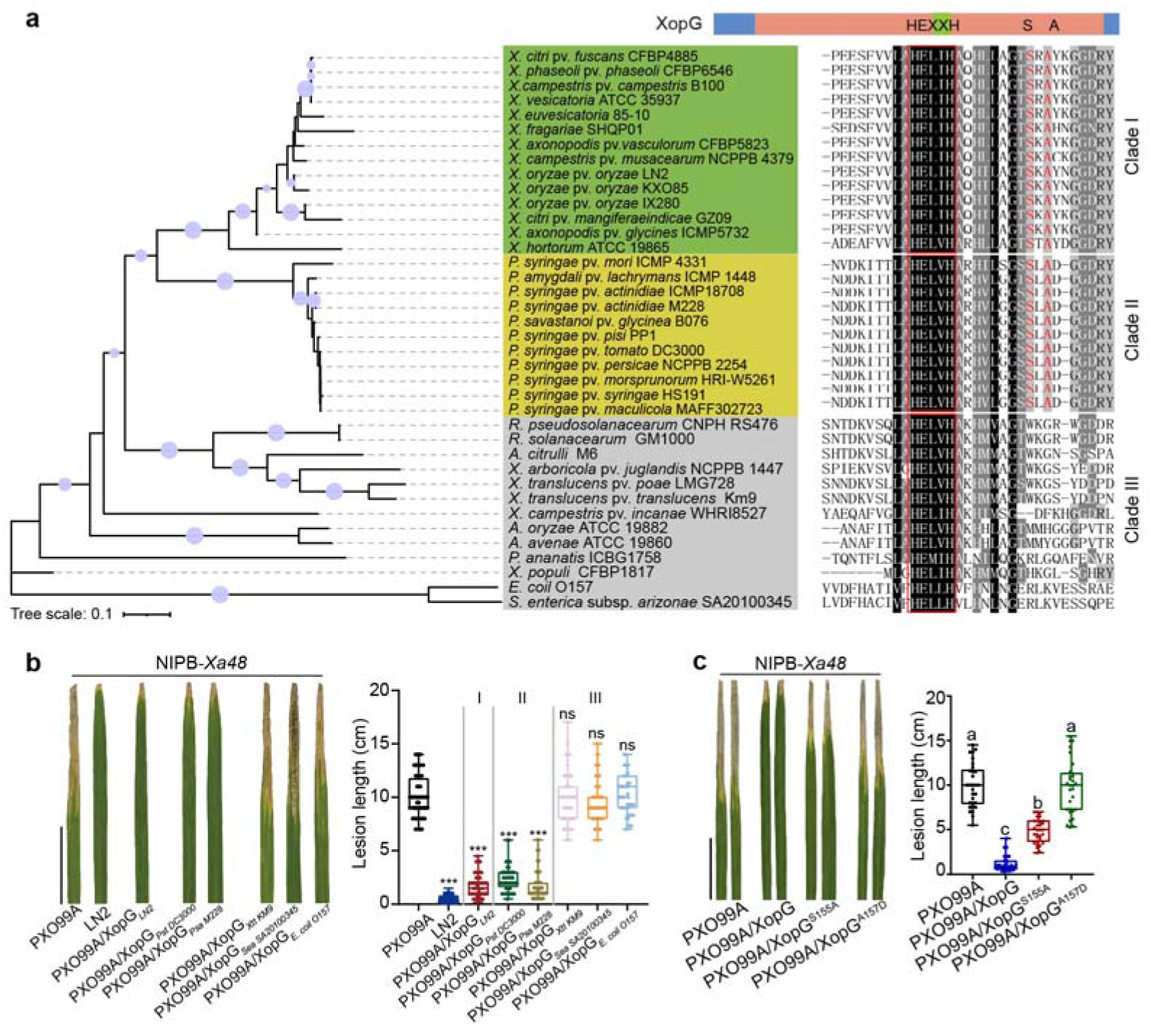
**Functionality and specificity analysis suggest XopG is a conserved effector in bacterial pathogens**. **a**, A phylogenetic tree and protein alignment of XopG and homologous proteins from different plant and animal bacterial pathogens were constructed using the Neighbor-Joining method, with 1,000 bootstrap replicates for statistical support, in MEGA11 software. Bootstrap values greater than 70% are indicated on the tree. XopG-like proteins are grouped into three clades. Note that the functional XopG *_LN2_*belongs to Class I, labeled with green (left). The conserved HEXXH motif is indicated with red box. Amino acids in red (XopG^S155^ and XopG^A157^) are conserved in Clade I and Clade II, but changed in Clade III (right). **b**, Disease resistance phenotypes of NIPB-*Xa48* inoculated with PXO99A/XopG*_LN2_*, PXO99A/XopG*_Pst DC3000_*, PXO99A/XopG*_Psa M228_*, PXO99A/XopG*_Xtt KM9_*, PXO99A/XopG*_Sea Serova_*, PXO99A/XopG*_E. coli O157_*, which contain three clades of XopG-like proteins. **c**, Disease resistance phenotypes of NIPB-*Xa48* inoculated with PXO99A expressing XopG^S155A^ and XopG^A157D^. Note that the S155 and A157 of XopG from Clade I and II were critical to its functional in XA48-mediated resistance. Data were shown as mean ± SD, *n* ≥ 20 (**b**, **c**), scale bars, 5 cm, asterisks represented statistical significance (\*\*\**P* < 0.001, two-tailed Student’s t-test). ns, not significant. Letters indicate significant differences (*P* < 0.05) determined by two-way analysis of variance (ANOVA) with Tukey’s test. Experiments were independently repeated twice with similar results (**b**, **c**).

**Extended Data Fig. 5.**
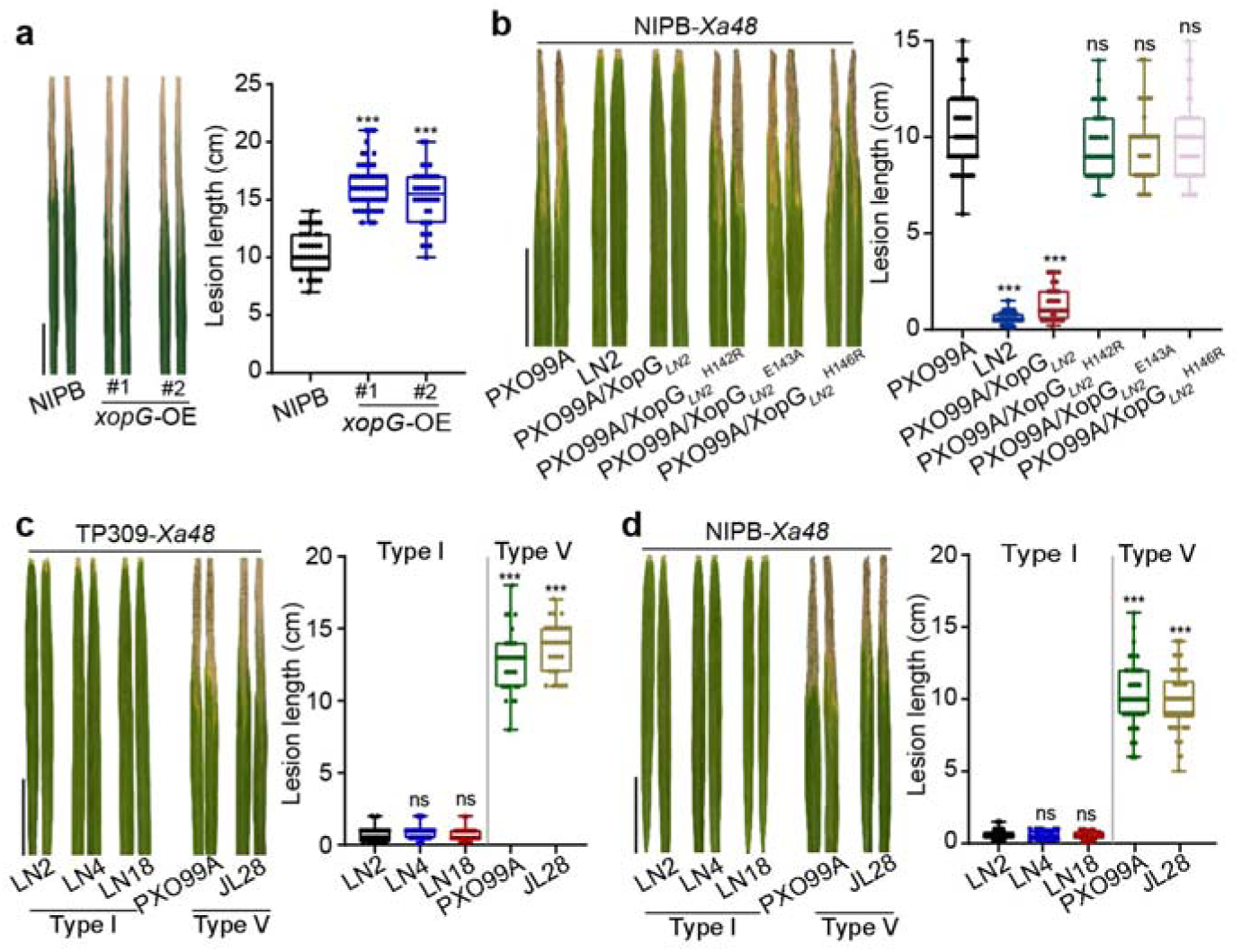
HEXXH domain of XopG contributes to trigger XA48-mediated resistance in rice. **a**, Decreased basal *Xoo* resistance of *xopG*-OE in NIPB. Leaves were inoculated with PXO99A. The lesion length was measured at 14 dpi. **b**, Disease resistance phenotype and lesion length of NIPB-*Xa48* inoculated with PXO99A/XopG*_LN2_*, PXO99A/XopG*_LN2_* ^H142R^, PXO99A/XopG*_LN2_* ^E143R^, and PXO99A/XopG*_LN2_* ^H146R^. LN2 and PXO99A-XopG*_LN2_* were used as avirulent controls. **c**, **d**, Avirulence and virulence detection of Type I and Type V of XopG variant strains on *Xa48*. Data were shown as mean ± SD, *n* ≥ 22, scale bars, 5 cm, asterisks represented statistical significance (\*\*\**P* < 0.001, two-tailed Student’s t-test). ns, not significant. Experiments were independently repeated twice with similar results.

**Extended Data Fig. 6.**
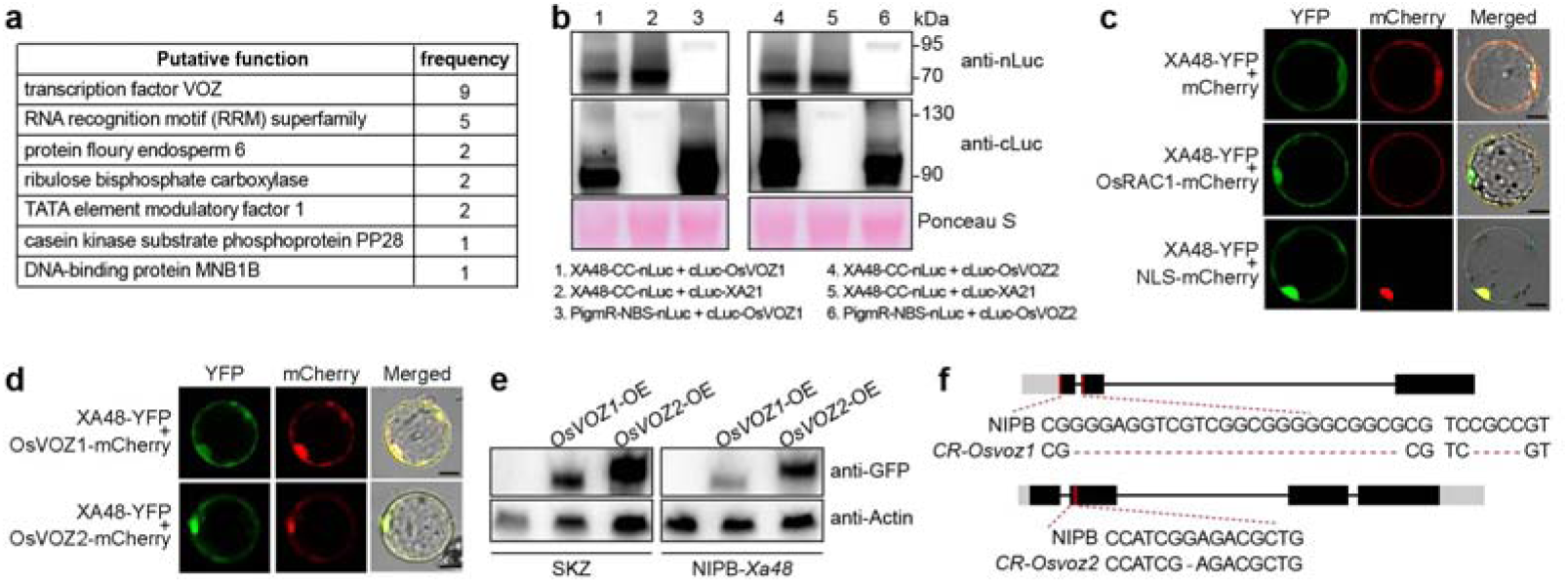
Identification of XA48-OsVOZ1/2 interaction. **a**, The candidate XA48-interacting proteins revealed by Y2H screen. Note that OsVOZ is the most hit protein. **b**, Western blots detected expression of XA48-CC-nLuc, cLuc-OsVOZ1 and cLuc-OsVOZ2 that were expressed in *N. benthamiana* for protein interaction. **c**, Representative images of XA48 subcellular localization. XA48-YFP was transiently expressed in rice protoplasts, which showed main localization to the PM/periphery and nucleus. OsRAC1-mCherry and NLS-mCherry served as a PM and nuclear marker, respectively. Scale bars, 10 µm. **d**, Co-localization of XA48 with OsVOZ1 and OsVOZ2 was detected in the PM and nucleus. Scale bars, 10 µm. **e**, Immunodetection of OsVOZs-GFP fusion protein in transgenic overexpression (OE) lines using anti-GFP antibody, with SKZ and NIPB-*Xa48* as wild type controls. Actin was detected as a loading control. **f**, Schematic of *OsVOZ1/2* knockout (KO) lines in NIPB background. Experiments were independently repeated twice with similar results (**b-e**).

**Extended Data Fig. 7.**
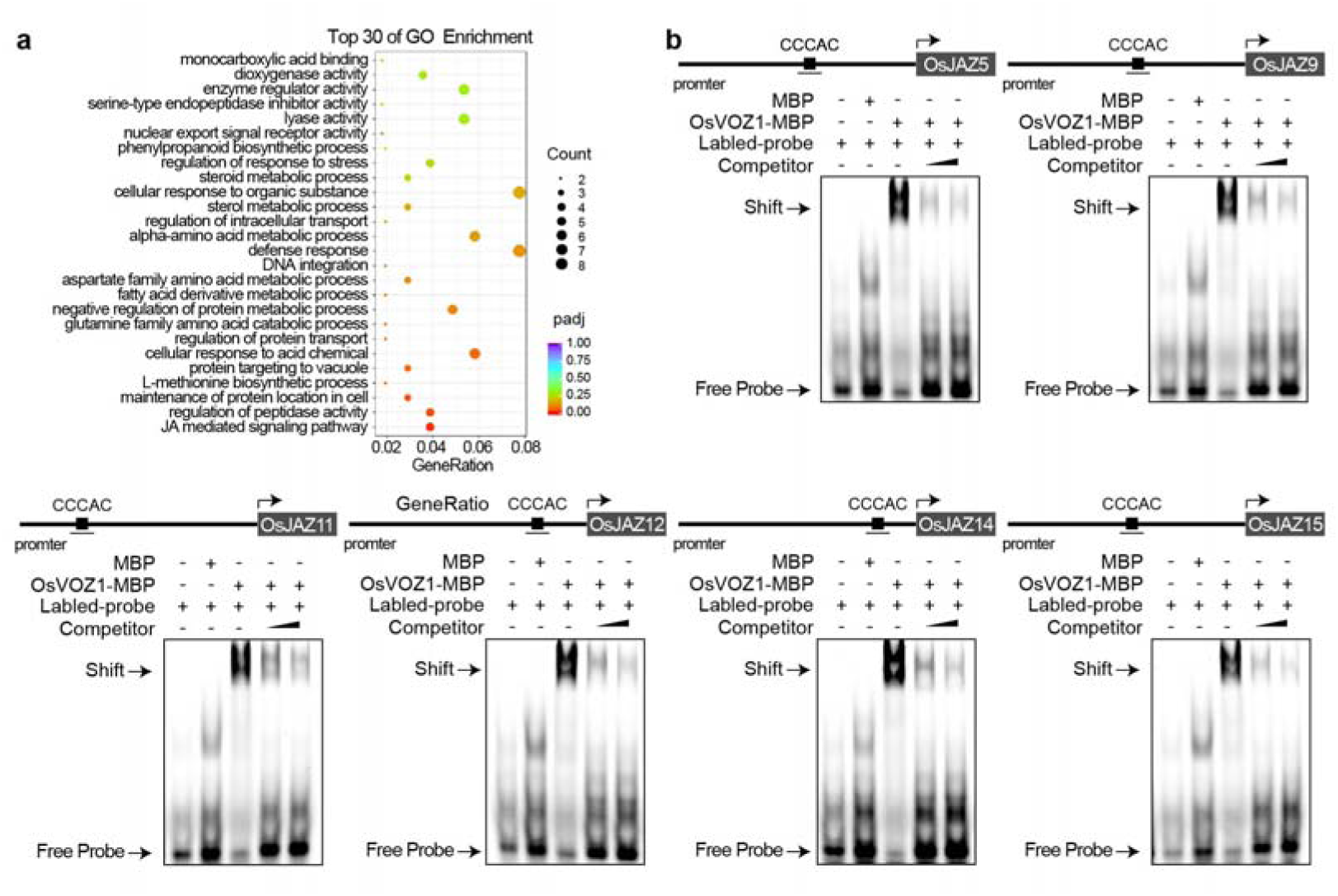
OsVOZ1 specifically binds to the promoter of *OsJAZ* family, which contain the conserved CCCAC motif. **a**, Gene Ontology (GO) analysis of differentially expressed genes in *CR-Osvoz1* and NIPB. Note that loss of *OsVOZ1* function activated genes involved in JA-related defense. **b**, EMSA was performed to investigate binding affinity of OsVOZ1 to the cis-elements in *OsJAZs* promoter. Biotin-labeled probes were incubated with MBP or OsVOZ1-MBP. Unlabeled competitor fragments were added to evaluate binding specificity. Experiments were independently repeated twice with similar results (**b**).

**Extended Data Fig. 8.**
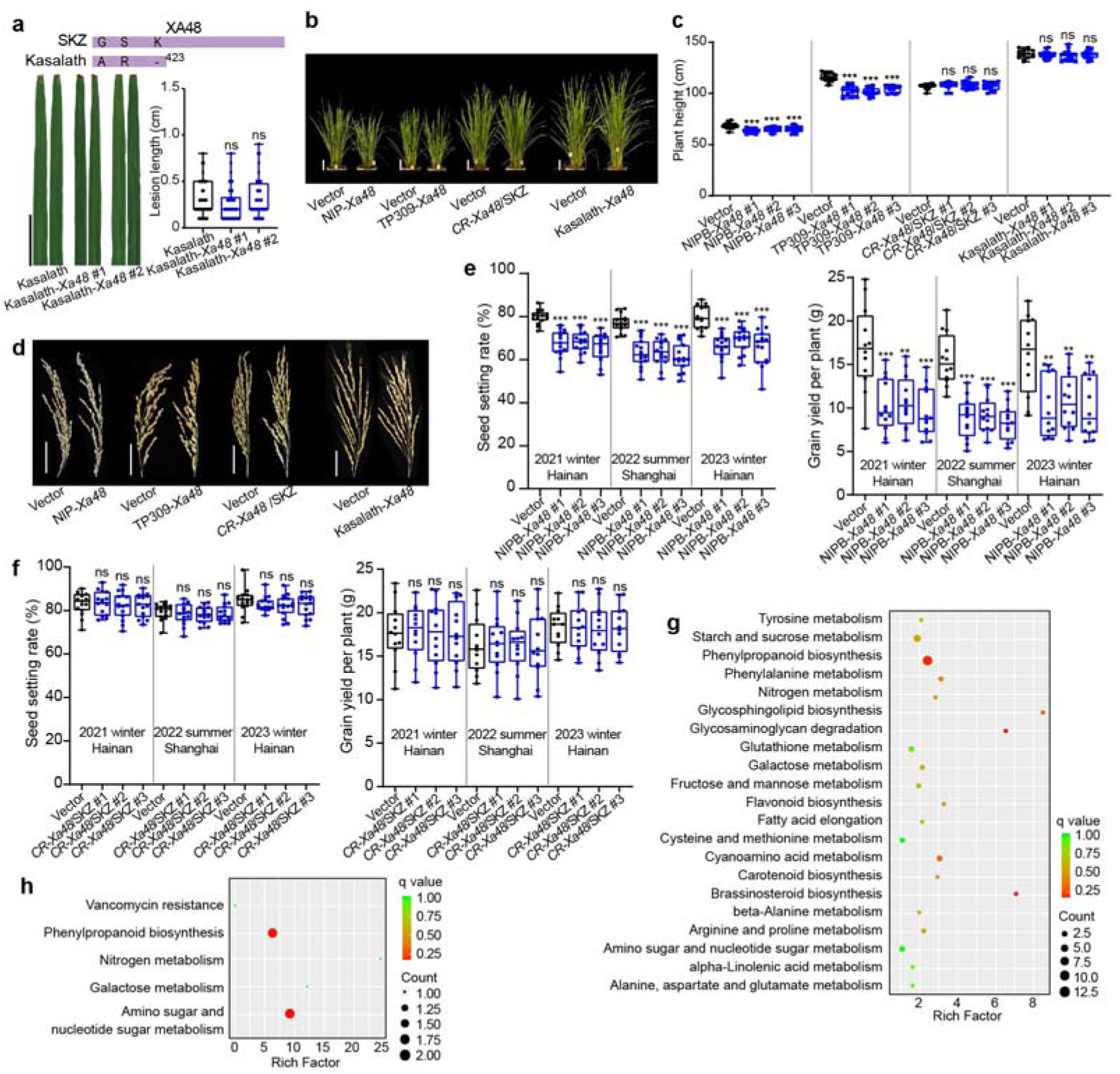
*Xa48* decreases grain yield in *japonica* but not in *indica*. **a**, Lesion length of *Xa48* complement lines in *indica* Kasalath that contains a truncated *Xa48* mutant (shown above) and harbors the *OsVOZ1^A^* allele. Note that wild type Kasalath is also resistant to J18, mediated by an additional unrecognized *Xa* genes. Scale bars, 5cm. **b-d**, Mature plants (**b**), plant height (**c**) and panicles (**d**) of NIPB-*Xa48*, TP309-*Xa48*, Kasalath-*Xa48*, and *CR-Xa48*/SKZ. Note that panicle size was not affected in these plants with or without *Xa48*, whereas *Xa48* led to a decrease in plant height in both *japonica* varieties NIPB and TP309, but not in *indica* rice SKZ and Kasalath. Scale bars, 10 cm (**b**) and 5 cm (**d**). **e**, The introduction of *Xa48* significantly decreased grain yield in *japonica* rice NIPB by reducing seed setting in multiple-location and season field trials during 2021, 2022 and 2023. **f**, Knockout of *Xa48* (CR-*Xa48*) did not affect grain productivity in *indica* rice SKZ in multiple-location and-season field trials during 2021, 2022 and 2023. **g**, KEGG analysis of differentially expressed genes in TP309-*Xa48* vs TP309 young panicle. Note that the *Xa48* introduction induced differential expression of many genes including those involved in sugar and amino acid metabolism, which may contribute to the seed development penalty in *japonica*. **h**, *Xa48* does not cause much difference of gene expression in *indica CR-Xa48*/*SKZ* vs *SKZ*. Data were shown as mean ± SD, *n* ≥ 15 (**a, c, e, f**), asterisks represented statistical significance (\*\*\**P* < 0.001, two-tailed Student’s t-test). ns, not significant. Experiments were independently repeated three times with similar results (**a, e, f**).

**Extended Data Fig. 9.**
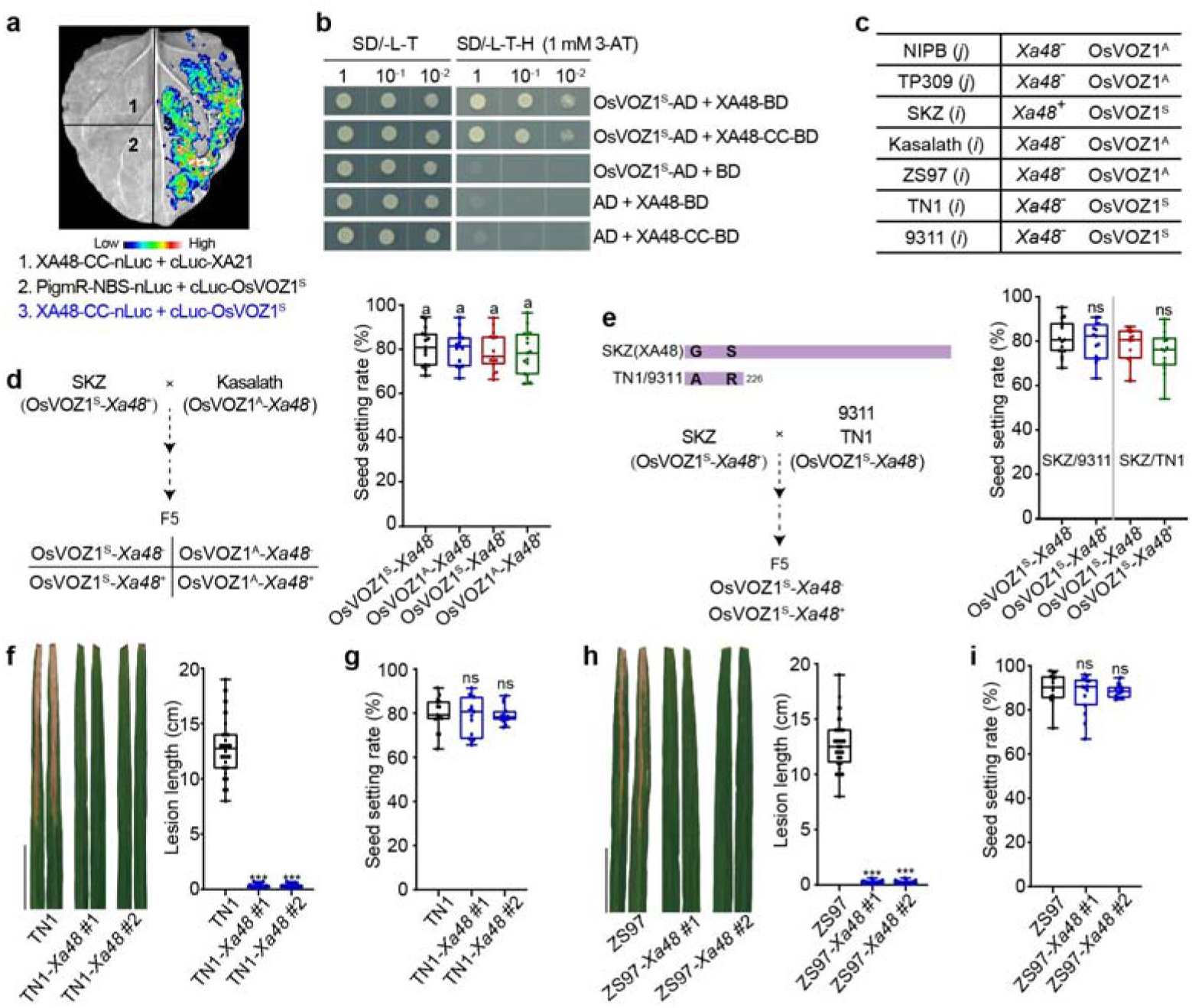
XA48-OsVOZ1^A/S^ immune modules shape different seed setting between *japonica* and *indica*. **a**, **b**, OsVOZ1^S^ interacts with XA48, determined by SLC (**a**) and Y2H (**b**) assays. **c**, The allelic variation of *Xa48* and OsVOZ1 in different varieties was shown in the *indica* (*i*) and *japonica* (*j*) varieties sued in the study. **d**, Development of *indica* inbreed lines with four combinations of OsVOZ1^A/S^ and *Xa48*, OsVOZ1^S^-*Xa48*^-^, OsVOZ1^S^-*Xa48*^+^, OsVOZ1^A^-*Xa48*^-^ and OsVOZ1^A^-*Xa48*^+^, derived from SKZ (OsVOZ1^S^) crossing to Kasalath (OsVOZ1^A^) (F_5_), which showed no difference in seed setting rate. **e**, Premature mutation of *Xa48* in *indica* TN1 and 9311 at codon position 226. SKZ crossing to 9311 and TN1, which brings the same OsVOZ1^S^ allele, to generate inbreed lines OsVOZ1^S^-*Xa48*^-^ and OsVOZ1^S^-*Xa48*^+^ (F_5_) respectively, which showed the same seed setting rate. **f-i**, Lesion lengths and seed setting rates of complement line TN1-*Xa48* (**f, g**) and ZS97-*Xa48* (**h, i**) (*indica*). The results indicated that the complement *indica* lines exhibited J18-resisatnce and no difference on seed setting rates. Scale bars, 5cm. Data were shown as mean ± SD, *n* ≥ 15 (**d-i**). For **d**, letters indicate significant differences (*P* < 0.05) determined by two-way analysis of variance (ANOVA) with Tukey’s test. For **e-i**, asterisks represented statistical significance (\*\*\**P* < 0.001, two-tailed Student’s t-test). ns, not significant. Experiments were independently repeated three times with similar results.

**Extended Data Fig. 10.**
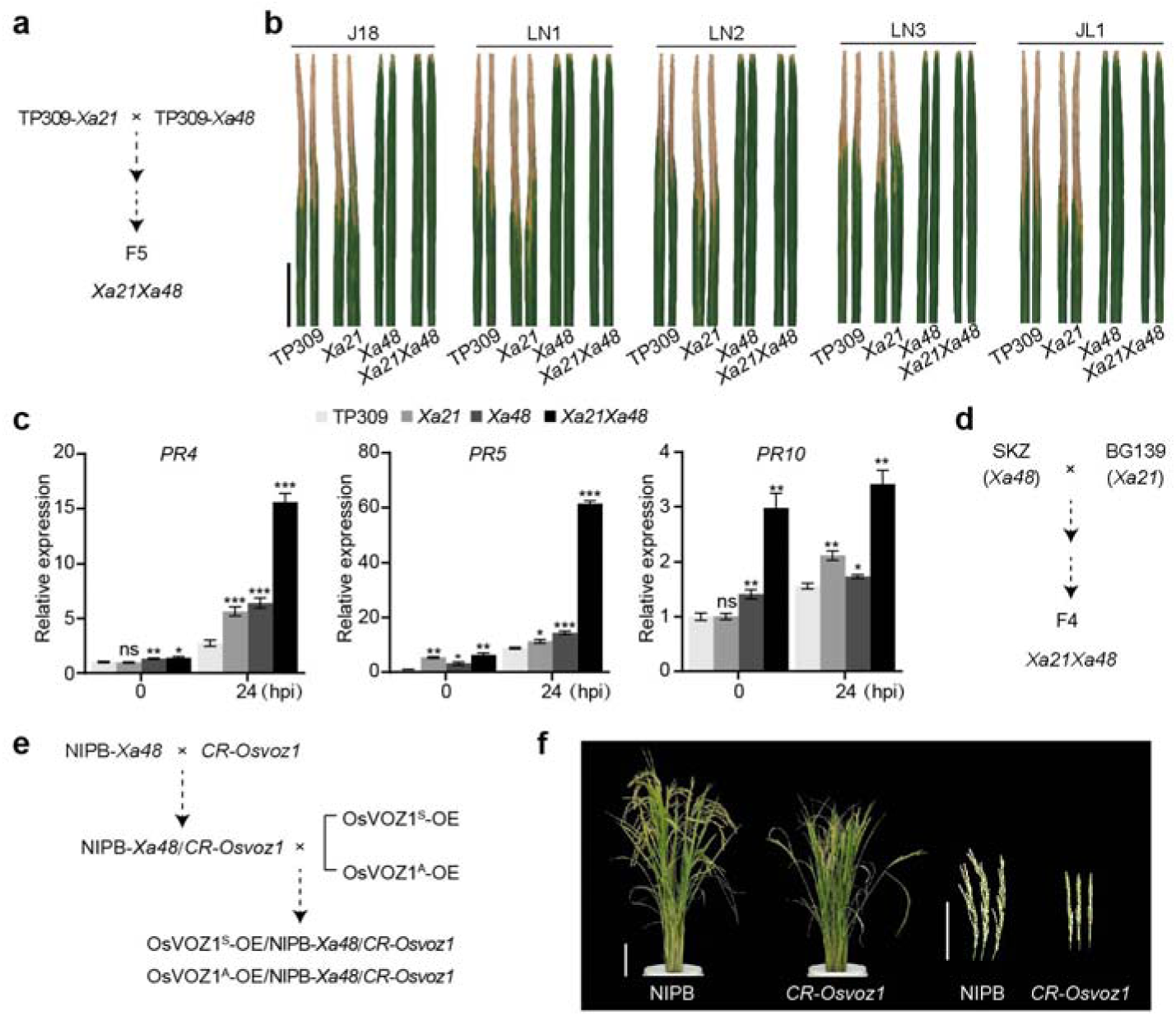
***Xa48*-*Xa21* stacking rice for broad-spectrum *Xoo* resistance was developed. a**, Development of transgenic TP309 (*japonica*) with integrating *Xa21Xa48* through crossing and selfing selection (F_5_ generation). **b**, Broad-spectrum disease resistance of *Xa21Xa48* plants against northeast Asian *Xoo* strain J18, LN1, LN2, LN3 and JL1. Scale bars, 5 cm. **c**, The defense genes *PR4*, *PR5*, and *PR10* expression were significantly enhanced in *Xa21Xa48* plants during *Xoo* infection. **d**, Development of *indica* rice combining endogenous *Xa48* and *Xa21* through crossing SKZ and BG139 (*Xa21*) and selfing (F_4_ generation). **e**, *De novo* development of *japonica* regaining *Xa48* and OsVOZ1^S^ by crossing *CR-Osvoz1*/NIPB and NIPB-*Xa48*. **f**, Mature plant and panicle of *CR-Osvoz1*, which showed defective growth and development phenotypes. Scale bars, 10 cm. Data were shown as mean ± SD, n = 3 (**c**), asterisks represented statistical significance (\**P* < 0.05, \*\**P* < 0.01, and \*\*\**P* < 0.001, two-tailed Student’s t-test). ns, not significant. Experiments were independently repeated twice with similar results (**c**).

